# Control of iron acquisition by multiple small RNAs unravels a new role for transcriptional terminator loops in gene regulation

**DOI:** 10.1101/2024.01.16.575843

**Authors:** Eugenio Solchaga Flores, Jonathan Jagodnik, Fanny Quenette, Alexey Korepanov, Maude Guillier

## Abstract

Small RNAs (sRNAs) controlling gene expression by imperfect base-pairing with mRNA(s) are widespread in bacteria and regulate multiple genes, including genes involved in iron homeostasis, through a wide variety of mechanisms. We previously showed that OmrA and OmrB sRNAs repress the synthesis of the *Escherichia coli* FepA receptor for iron-enterobactin complexes. We now report that five additional sRNAs, namely RprA, RybB, ArrS, RseX and SdsR, that respond to different environmental cues, also repress *fepA,* independently of one another. While RprA follows the canonical mechanism of pairing with the translation initiation region, repression by ArrS or RseX requires a secondary structure far upstream within the long *fepA* 5’UTR. We also demonstrate a dual action of SdsR, whose 5’ end pairs with the *fepA* translation initiation region while its 3’ end behaves like ArrS or RseX. Strikingly, mutation analysis shows a key role for the loops of these sRNAs intrinsic terminators in the regulation. Regulation furthermore depends on both the Hfq chaperone and the RNase E endonuclease. Overall, our data strongly suggest that FepA levels must be tightly controlled under a variety of conditions, and highlight the diversity of mechanisms that underly the regulation of gene expression by sRNAs in bacteria.

## INTRODUCTION

Bacteria have the ability to adapt to a wide diversity of environments by regulating their gene expression in response to a multitude of internal and external signals. This genetic control relies on protein and RNA regulators that can target the different stages of gene expression. Among these, small RNAs (sRNAs) acting as post-transcriptional regulators via imperfect pairing to target mRNAs have been extensively studied. They exist in virtually all bacteria, and studies in model organisms revealed some key aspects of their biological functions and modes of action (reviewed in (1)). Their pairing to target mRNA(s) typically affects the translation and/or the stability of the mRNA, either positively or negatively. In some cases, often species-dependent, their action requires protein chaperones such as Hfq or ProQ. Importantly, sRNAs can be primary transcripts or processing products originating from an mRNA (or even from a longer isoform of an sRNA) and they often respond to environmental changes because their synthesis, or that of their precursors, is controlled by transcription factors. As a result, bacterial gene expression is controlled via complex regulatory circuits integrating transcriptional and post-transcriptional controls, often mediated by proteins and sRNAs, respectively. In addition, the regulation of transcription factors synthesis by sRNAs, often in a direct or indirect feedback control pathway, may also have major implications for bacterial gene expression in response to changing environment (2).

A single sRNA typically targets multiple mRNAs, which is facilitated by imperfect base-pairing interactions, and thereby controls the expression of many genes. Hence, sRNAs can be involved in multiple pathways, ranging from central carbon metabolism to membrane stress responses. Among those processes, metal homeostasis, and in particular iron homeostasis, has been repeatedly found to be controlled by sRNAs in a wide range of bacterial species, sometimes in conjunction with the Fur transcriptional repressor (3–5). In *Escherichia coli* (*E. coli*) for instance, the Fur-controlled RyhB sRNA was shown early on to repress several genes encoding iron storage or iron-containing proteins (6). These regulations not only expand the Fur regulon, but also result in indirect gene activation by the Fur repressor. RyhB also activates the synthesis of enterobactin (7), the most potent iron siderophore. In addition, later studies demonstrated a role for RyhB and other sRNAs in controlling the synthesis of several iron-siderophore receptors, such as CirA, Fiu, FecA or FepA (8–11). Of note, *fepA* encodes the receptor for iron-enterobactin complexes, and therefore plays a key role in iron acquisition. This sRNA control of metal homeostasis is likely important for bacterial virulence, as both metal sequestration and intoxication can be encountered by bacteria during host infection (12).

From a mechanistic standpoint, control by sRNAs most often relies on their pairing to the translation initiation region (TIR) of the target-mRNAs. This interaction impedes, or more rarely favors, the binding of the 30S ribosomal subunit and therefore impacts translation initiation. However, this canonical mechanism is far from being the only one and many others have already been described. One illustration of sRNAs pairing outside the TIR was provided by the regulation of *fepA* by the sRNAs OmrA and OmrB. These two partly redundant sRNAs pair to the coding sequence of *fepA*, more than 20 nucleotides (nts) downstream of the start codon, and impede the formation of a secondary structure that activates translation initiation (10). Consistent with this model, the long *fepA* 5’UTR was shown to be mostly dispensable for control by OmrA and OmrB.

Intriguingly however, other mRNAs with long 5’UTRs, such as *flhDC* or *csgD*, are known to be regulated by multiple sRNAs (13–20). Similarly, the >140-nt *fepA* 5’UTR could thus serve as a hub for post-transcriptional regulation by actors other than OmrA or OmrB. This prompted us to search for additional sRNAs regulating *fepA*. Our results show that at least seven sRNAs repress *fepA* expression, and their action depends on different regions of the *fepA* mRNA. Furthermore, in at least two cases, the regulation relies on the terminator loop of the sRNAs rather than on the previously described base-pairing regions. Overall, this study reveals that sRNA regulation can act in yet undescribed ways and unravels new connections between sRNAs and iron uptake.

## MATERIAL AND METHODS

### General microbiology techniques

*E. coli* strains used in this study are derivatives of the MG1655 wt strain and are listed in Table S1; the only exception is the strain NEB5alpha (NEB reference #c2992) used for cloning purposes. Unless otherwise indicated, they were grown in LB medium, supplemented with antibiotics when needed at the following concentrations: Ampicillin 150 μg/ml, Tetracycline 10 μg/ml, Chloramphenicol 10 μg/ml; Kanamycin 25 μg/ml.

Deletion mutants of *fepA* or of the different sRNAs were constructed by recombineering to replace the gene of interest by an antibiotic resistance cassette. Some resistance cassettes were surrounded by FRT sites and, when needed, were eliminated by Flp-FRT recombination using the pCP20 plasmid as previously described (21).

Reporter fusions were constructed by recombineering into strain MG1508 (for *lacZ* fusions, see (22)) or MG2352 (for *mScarlet* fusions, see (21)). A single mutation in the -35 region of the P_tet_ promoter (T<u>T</u>GACA->T<u>G</u>GACA, referred to as <u>low</u> P_tet_, (21)) was introduced in some cases to lower transcription.

Plasmids overproducing different sRNAs derive from the pBRplac empty vector (9). Most plasmids used in Fig. 1A are from (23), except for pCpxQ and pDapZ (24), and pMcaS, pRaiZ, pNarS, pCsrC, pSgrS and pArrS that were constructed in this work (Table S1), either because they were not present in (23) or because the way they were constructed did not allow resistance to both ampicillin and tetracycline. For the new plasmids, the overproduction of the sRNAs was verified by Northern-blot (Fig. S1). Plasmids were constructed using the NEB HiFi DNA assembly master mix following manufacturer’s instructions. When needed, pBRplac derivatives were mutagenized using PCR with mutagenic primers and the Pfu Turbo DNA polymerase (Agilent), followed by DpnI digestion of the template, and transformation into NEB5alpha cells. Plasmid construction was systematically checked by sequencing. In general, ampicillin was used to maintain the plasmids, and 100 μM IPTG was used to induce the sRNAs transcription; for the fluorescence experiments where cells were grown for 16 hours, tetracyclin was used instead for plasmid maintenance and 250 μM IPTG to induce transcription of the sRNAs. Primers used for strains and plasmids construction are listed in Table S2.

**Figure 1.**
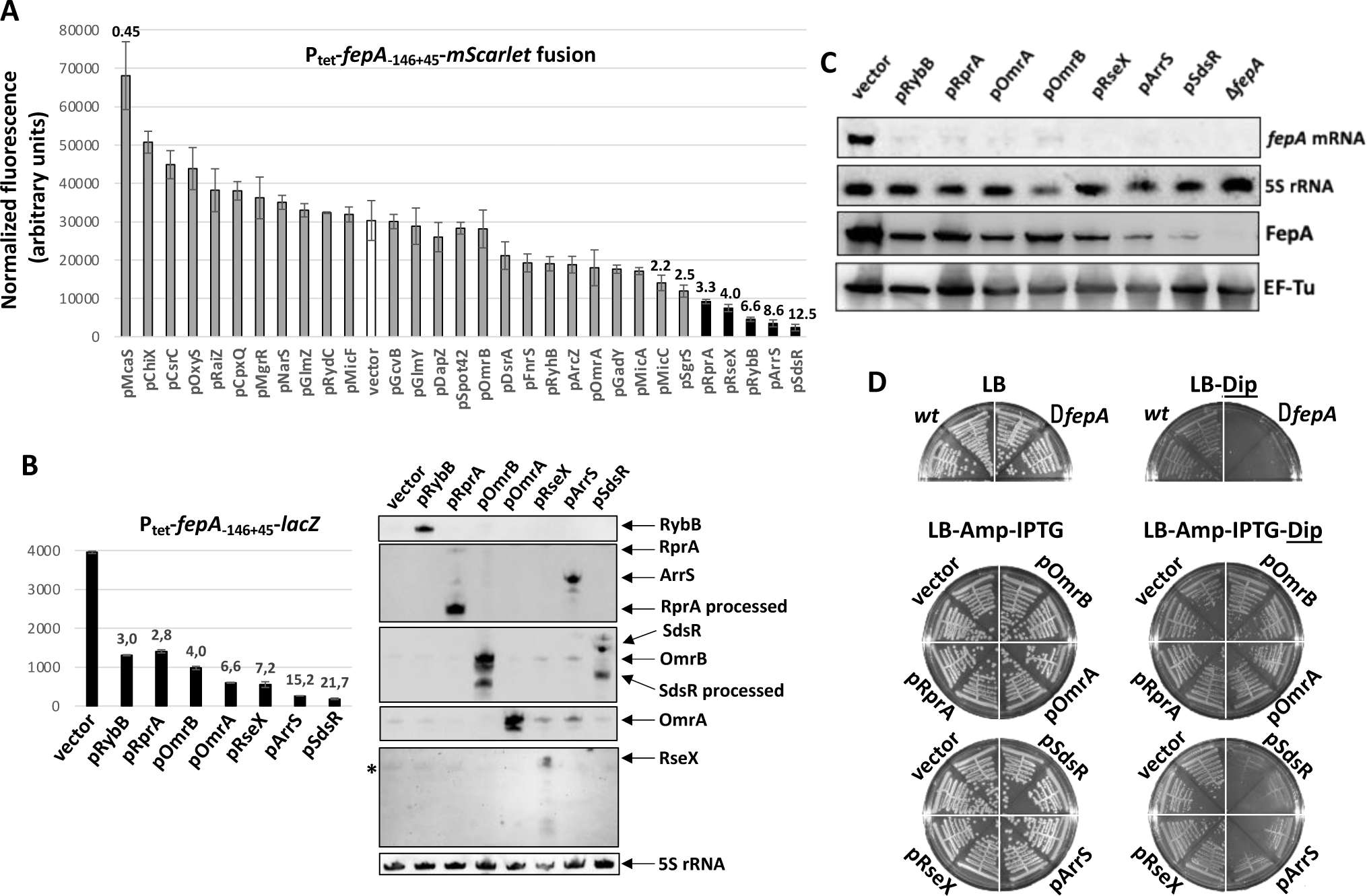
*E. coli fepA* expression is repressed by RprA, RybB, OmrA, OmrB, ArrS, RseX, and SdsR sRNAs. (A) Fluorescence of the P_tet_-*fepA-mSc* reporter in strain FQ128 was measured in the presence of plasmids overproducing 30 different sRNAs, mostly Hfq-dependent. Shown is the fluorescence normalized to the absorbance at 600nm in exponential phase (arbitrary units). Numbers above the bars indicate the repression factors compared to the empty vector (white bar) when the plasmid has an effect greater than 2-fold, and the black bars show the plasmids that have an effect greater than 3-fold. (B) The specific β-galactosidase activity of the P_tet_-*fepA_-146+45_-lacZ* fusion in strain JJ389 was measured in the presence of plasmids overproducing the indicated sRNAs. The sRNAs levels were analyzed by Northern blot in parallel, using the detection of the 5S rRNA as a loading control. (C) The *fepA* mRNA and FepA protein levels were analyzed by Northern and Western blot, respectively, from the *Δfur* strain ES021 transformed by the indicated plasmids and grown to exponential phase. The levels of the 5S rRNA, and of the EF-Tu protein, were used as loading controls. (D) The growth of *fepA^+^* or *ΔfepA* strains (MG1325 and ES024, respectively) was followed on LB plates supplemented or not with 250 μM 2-2’-dipyridyl (dip.). And the growth of strain JJ389 transformed by the indicated plasmids was followed on LB-Amp-IPTG plates supplemented or not with 250 μM dip. Plates were incubated at 37°C overnight, or for 48 hours when containing dip.

### Fluorescence measurements

The fluorescence of *mScarlet* fusions was assessed as previously (21). Briefly, cells were grown in minimal A medium supplemented with 0.25% casamino acids, 0.5% glycerol and 1 mM MgSO_4_ (CAG) with tetracycline overnight, diluted 200-fold in fresh medium supplemented with 250 μM IPTG, and optical density at 600nm (OD_600_) and fluorescence using an excitation wavelength of 560nm and an emission wavelength of 600 nm with a bandwidth of 15 nm were followed for 16 hours. The value of fluorescence at the point where OD_600_ is closest to 0.4 was used to calculate the normalized fluorescence (=fluorescence/OD_600_) for each culture. Values of fluorescence shown in the figures correspond to the average normalized fluorescence of three independent replicates (2 replicates for the screen of Fig. 1A) and the error bars indicate the standard deviations. For the screen of Fig. 1A, strains transformed with pDicF, pCyaR, pIS118, pMicL or pSdsN_137_ did not grow enough in 96-wells plate to allow fluorescence measurement.

### β-galactosidase assays

Strains were grown in LB medium containing antibiotics and IPTG as needed. Miller assays were performed as previously described (25) using chloroform and SDS to lyse the cells. For the experiments with a large number of samples (Fig. 5 and S4), cells were diluted 200-fold from saturated overnight cultures in 200 μl medium in 96-wells plates and grown with shaking for 4.5 hours. After OD_600_ was measured, 100 μl of cells were mixed with 50 μl lysis buffer containing polymyxin B and DTT (as in (26)), incubated at room temperature for 30 minutes, and 50 μl ONPG 4mg/ml was added. Samples were covered with 50 μl mineral oil and incubated at 28°C. The absorbance at 420nm and 550nm was followed and the β-galactosidase was calculated with the Miller formula after 75 minutes incubation. All β-galactosidase data are shown as average of three independent replicates with error bars indicating standard deviations.

### RNA extraction, Northern blots and primer extension

RNA was extracted using the hot-phenol method mostly as in (9). 650 μl of cell cultures in LB were directly mixed with phenol, while 2 ml of cultures in CAG were first centrifuged at 4°C and the cell pellets resuspended in 650 μl PBS1X prior to be mixed with phenol. After phenol extractions, RNA was ethanol-precipitated and equal amounts of total RNA were separated by electrophoresis. For the analysis of sRNAs or mRNAs, 3 to 8 μg of total RNA were loaded on denaturing acrylamide gels, or 5 μg RNA were loaded 5 on agarose gels, respectively. After migration, RNAs were transferred to Hybond-N+ membranes, and specific RNAs were detected using specific biotinylated probes and the same detection procedure described in (24).

For analysis by primer extension, 5 μg of total RNA were mixed with 2 pmol Cy5-labeled probe, denatured at 80°C for 3 minutes, flash-frozen in dry ice-cold ethanol and thawed on ice. After 5 min-incubation at 37°C, primer extension was performed with Superscript III following manufacturer’s instructions. cDNAs were separated on a denaturing 6% polyacrylamide sequencing gel and visualized with a Typhoon fluorescent scanner set up for Cy5 detection.

### Western blot

Protein samples preparation and western blotting were mostly as described previously (22), with proteins being separated on Mini-protean TGX precast gels 4-15% (Biorad). Transfer was done with the Transblot Turbo RTA transfer kit Nitrocellulose and detection was performed with the Clarity Max Western substrate (Biorad) following manufacturer’s instructions. The anti-FepA is a kind gift from Kathleen Postle (27) and was used overnight at a 1:5000 dilution in PBS-Tween 0.01% with 2.5% milk at 4°C; the antibody against EF-Tu was used at a dilution of 1:10000 in the same solution.

### In vitro transcription and RNA structure probing

*In vitro* RNA synthesis was done with the T7 RNA polymerase using PCR products as templates and the T7 Megascript kit (Ambion) following manufacturer’s instructions. 2 or 3 G were added downstream of the T7 promoter to enhance efficiency of transcription for *fepA*, but they were omitted for the sRNAs.

Probing using DMS or CMCT was done as described in (10). For lead-acetate probing, the mRNA-sRNA mixes were denatured at 80°C for 3 minutes and flash-frozen in dry ice-cold ethanol. They were equilibrated in a “structure buffer”(final composition is 10mM Tris-HCl pH 7,5; 300mM NaCl; 5mM EDTA pH 7,5 and 10mM MgCl_2_) after denaturation, and incubated at 37°C for 10 min. Lead-acetate was then added at the final concentration of 5mM and incubation continued for 3 min at 37°C. Reactions were stopped with 50mM EDTA, and RNA were ethanol-precipitated with 2 μl glycoblue. After resuspension in water, RNA was mixed with 2 pmol Cy5-labeled probe and subjected to primer extension as described above.

## RESULTS

### At least seven sRNAs repress the expression of *fepA* in *E. coli*

In order to identify other putative sRNA regulators of *fepA*, we took advantage of a plasmid library developed by Mandin and Gottesman (23) allowing the overproduction of multiple Hfq-dependent sRNAs to which we added plasmids overproducing the more recently described sRNAs McaS (19), ArrS (28), CpxQ (29), DapZ (30), NarS (31) as well as the ProQ-dependent RaiZ (32) and the CsrA-binding CsrC (33) sRNAs (Fig. S1). To study the effect of these sRNAs on *fepA* expression, the fluorescence of a *fepA-mScarlet-I (fepA-mSc)* translational reporter fusion was measured after transformation by these 30 different plasmids. This *fepA-mSc* construct carries the nts -146 to +45 of *fepA* mRNA relative to its start codon fused in frame with the *mSc* coding sequence, so that the 5’-end of the *fepA-mSc* mRNA corresponds to the predominant 5’-end of *fepA* detected by primer extension in exponentially growing cells (Fig. S2A). The fusion is furthermore expressed from a P_LtetO-1_ promoter (hereafter referred to as P_tet_, (34)) and is thus independent of control exerted on the *fepA* promoter, such as repression by Fur.

In this experiment, 5 out of the 30 sRNAs tested affected the fluorescence of *fepA-mSc* more than 3-fold, always negatively: RprA, RseX, RybB, ArrS and SdsR (Fig. 1A). Surprisingly, the sRNAs OmrA and OmrB that target a structure in the early coding sequence of *fepA* mRNA (10) had no significant effect in this experiment, most likely because this *fepA-mSc* construct is not a suitable reporter in this case (see Fig. S2 for details).

To validate these five novel regulators of *fepA*, we then looked at their effect on the expression of both a *fepA-lacZ* translational fusion (again transcribed from the P_tet_ promoter), and the *fepA* endogenous gene. All five plasmids, as well as pOmrA and pOmrB, efficiently repressed the expression of *fepA-lacZ*, decreasing the β-galactosidase activity from 2.8-to more than 20-fold (Fig. 1B) and induced a clear decrease in *fepA* mRNA and FepA protein levels (Fig. 1C). This control of *fepA-lacZ* and endogenous *fepA* by OmrA and OmrB confirms that the lack of regulation of *fepA-mSc* by these sRNAs is context-dependent, and that *fepA-mSc* is not a good reporter for regulations targeting the translation-activating stem-loop. Overall, these results show that at least seven sRNAs repress *fepA* expression.

As FepA is the receptor for iron-enterobactin complexes, it plays a major role in iron acquisition and *fepA*-deleted cells display a severe growth defect in iron-depleted conditions, e.g., in the presence of an iron chelator such as 2-2’-dipyridyl (dip, Fig. 1D). We also tested the effect of some of the sRNAs repressing *fepA* under these conditions and found that overproduction of either SdsR and ArrS inhibited growth of iron-starved cells (Fig. 1D). This is consistent with the observation that these two sRNAs display the strongest *fepA* repression in several experiments (Fig. 1, panels A-C).

### Independent action of the 7 sRNAs, and of Hfq, to repress *fepA*

We next checked whether the effect of some of these seven sRNAs could be mediated by one (or more) of the other six sRNAs, or whether they act independently to repress *fepA*. To address this question, we analyzed the control of *fepA* by the different plasmids in cells deleted for the chromosomal copies of the *omrA*, *omrB*, *rprA*, *rybB*, *arrS*, *sdsR* and *rseX* genes (Δ7 strain, Fig. 2A). This was done using the translational *fepA-lacZ* fusion as a reporter, transcribed here from a modified version of the P_tet_ promoter (lowP_tet_) to decrease its expression and potentially be able to detect up-effects of the sRNAs deletions. Repression of *fepA* by all plasmids was similar in the wt and Δ7 strains, clearly showing that the action of each of these seven sRNAs is independent of the six others (Fig. 2A).

**Figure 2.**
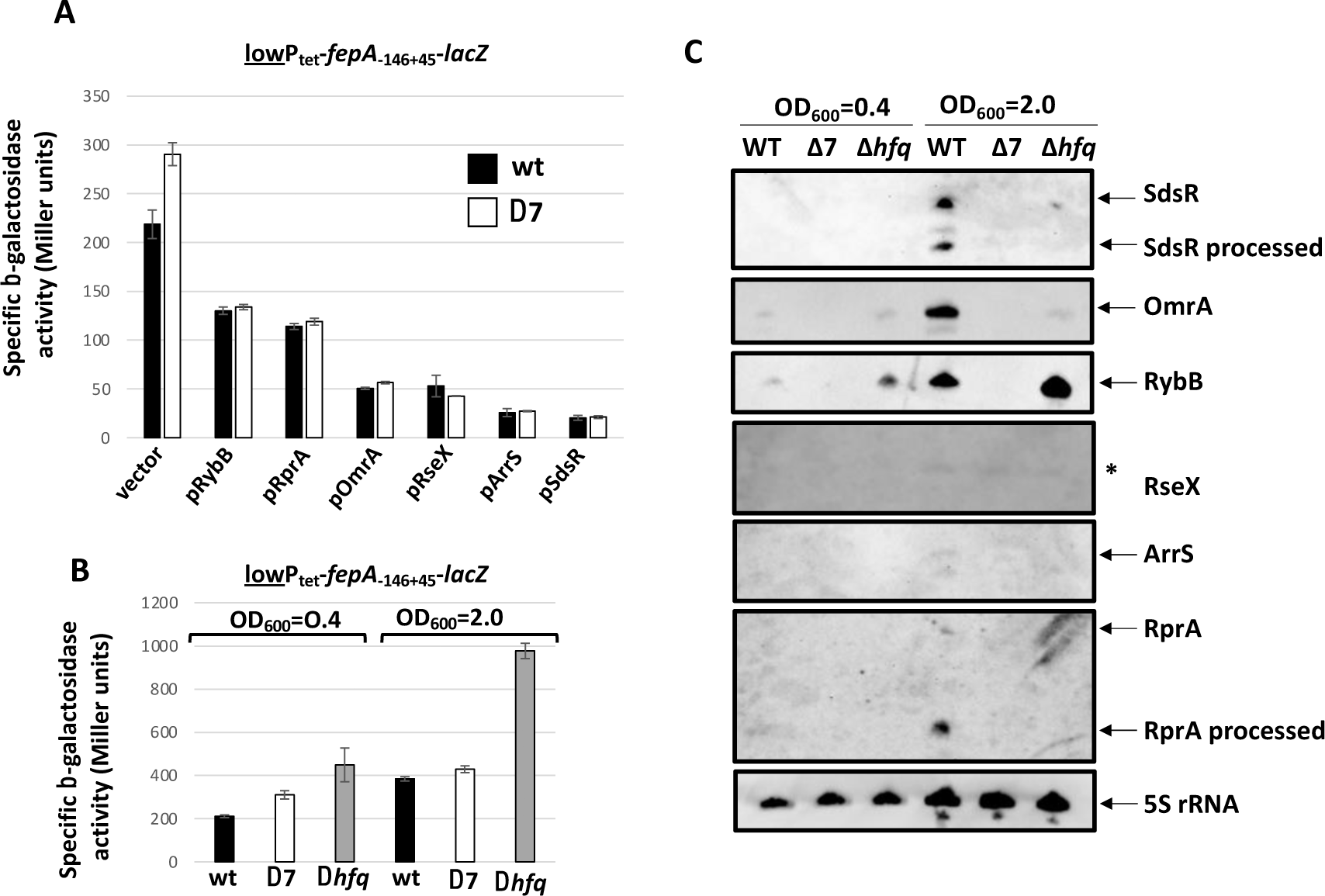
The 7 sRNAs, and most likely Hfq, independently repress *fepA*. (A) The specific β-galactosidase activity of the lowP_tet_-*fepA_-146+45_-lacZ* fusion was measured in the presence of plasmids overproducing the indicated sRNAs in strains that are wt (OK410) or deleted for *omrAB*, *rprA*, *rybB*, *sdsR*, *arrS* and *rseX* (Δ7, ES250). (B) The activity of the same fusion was measured in the wt, Δ7 or Δ*hfq* background (strains OK410, ES250 and ES234) in exponential or early stationary growth phase, i.e., at an absorbance at 600nm of 0.4 or 2, respectively. (C) Total RNA was extracted from the cultures used in panel B and the indicated sRNAs analyzed by Northern blot. The asterisk on the Northern probed for RseX indicates a faint signal due to cross-hybridization as it is visible in all samples, including the Δ7.

Interestingly, this experiment also shows that the deletion of the seven sRNAs up-regulated *fepA* expression by a factor of about 1.4-fold (compare wt and Δ7 in presence of the vector control). This difference was similar in the absence of plasmids in exponential phase (1.5-fold increase in the Δ7 strain at OD_600_=0.4; Fig. 2B), but was no longer visible at the onset of stationary phase (OD_600_=2, Fig. 2B). This is surprising as Northern blot analysis shows that at least four of these sRNAs, namely SdsR, RprA, OmrA, and RybB, are more abundant at OD_600_=2 than OD_600_=0.4, consistent with published data (e.g. (35–37), also suggesting that this is true for OmrB), while the other two sRNAs, RseX and ArrS, were not or barely detected by Northern blot in either condition (Fig. 2C). It is possible that (an)other factor(s) are involved in repressing *fepA* expression in the Δ7 strain. Lastly, this experiment showed that the deletion of *hfq* had a greater impact than deletion of the seven sRNAs, since *fepA* expression was increased in the *Δhfq* strain by 2.2- and 2.5-fold at OD=0.4 and OD=2, respectively (Fig. 2C). These data strongly suggest that Hfq also plays an independent role in *fepA* repression in addition to the repression by the 7 sRNAs.

### RprA and SdsR repress *fepA* via direct binding to the TIR

To identify possible mechanisms of regulation by the five newly identified *fepA* repressors, we searched for potential base-pairing interactions between RprA, RybB, RseX, ArrS or SdsR sRNAs and the (-146+45) region of the *fepA* mRNA using IntaRNA 2.0 (38). The most convincing predictions were found for the RprA and SdsR sRNAs, and the pairing is predicted with the *fepA* TIR in both cases (Fig. 3). We tested these putative interactions by introducing compensatory changes in either the sRNA and/or the *fepA-lacZ* fusion. In the case of RprA, mutating either the sRNA or the mRNA strongly impaired regulation, while restoring the interaction by mutating both restored control (2.5- and 2.1-fold repression in the wt/wt and mutant/mutant context, respectively; Fig. 3A). With SdsR, control of the *fepA_-146+45_-lacZ* fusion was also more efficient in the wt/wt (36-fold repression) or mutant/mutant (24-fold) context than when pairing was disrupted with mutation in either SdsR or in *fepA-lacZ* (Fig. 3B). These data support the predicted interactions of RprA or SdsR to the *fepA* TIR that presumably compete with 30S binding and thus inhibit translation initiation. For SdsR, however, repression is still visible in the presence of these mutations that are expected to disrupt the SdsR-*fepA* pairing (12-fold repression of wt *fepA-lacZ* by SdsRmutβ, and 15-fold for *fepAmutβ-lacZ* by wt SdsR, Fig. 3B). This suggests that the pairing of SdsR to *fepA* Shine-Dalgarno (SD) region only explains part of the observed regulation, and that a second action of SdsR, involving another mechanism, contributes to the full regulation. This could explain the strong effect of SdsR on *fepA* expression, which is rather unusual for an sRNA, even when overproduced.

**Figure 3.**
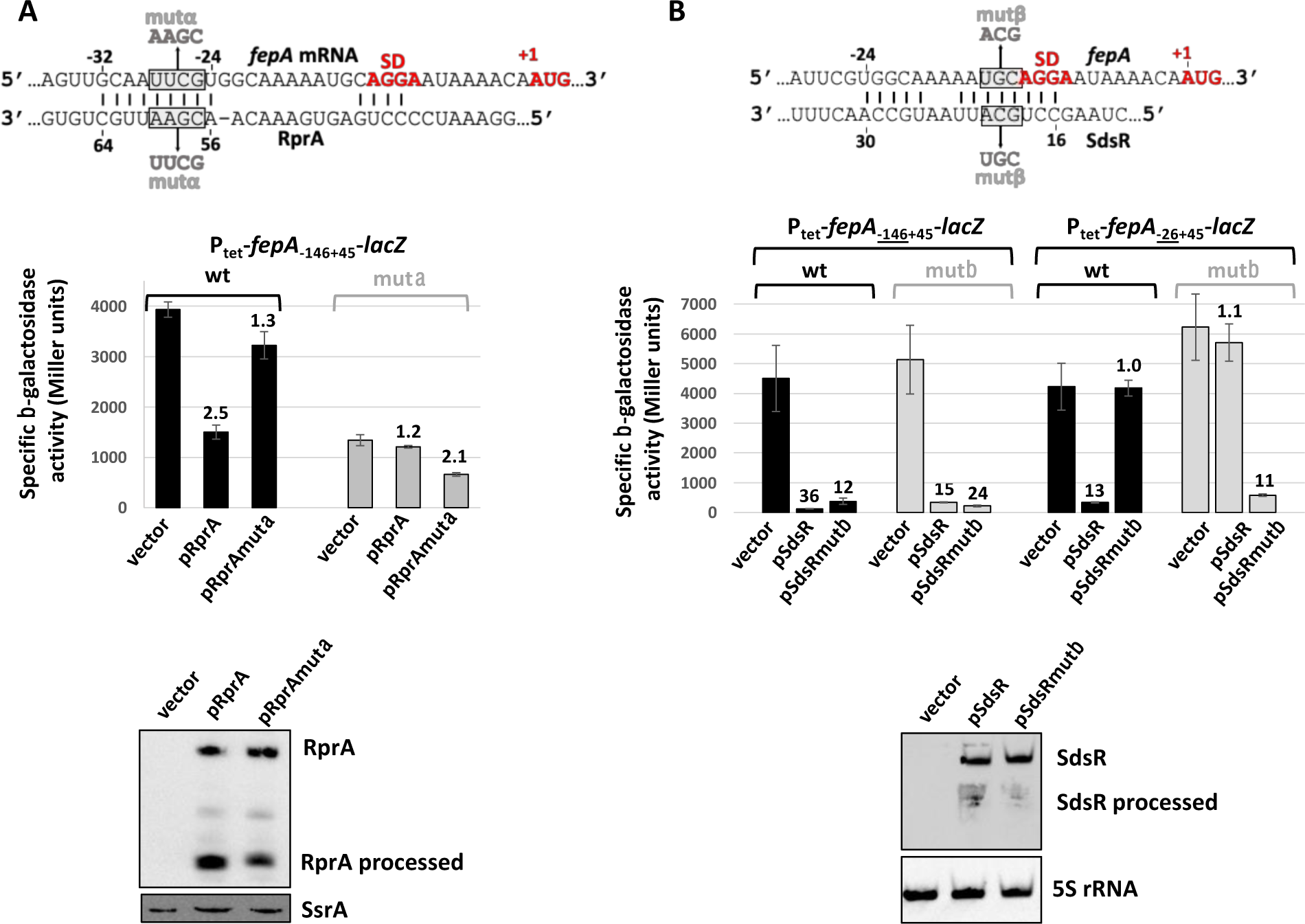
Pairing of RprA and SdsR sRNAs to the *fepA* TIR. Pairing predictions between the *fepA* mRNA and the RprA (A) or SdsR (B) sRNA are shown with the mutations introduced in the P_tet_-*fepA-lacZ* fusions and/or the sRNA overexpressing plasmid. The β-galactosidase activity of fusions carrying wt or mutant *fepA* region (-146+45) (panels A and B) or (-26+45) (panel B only) was measured in the presence of the indicated plasmids. The numbers above the bars give the repression factors compared to the empty vector. The levels of the wt or mutant RprA (A) or SdsR (B) sRNA were analyzed by Northern blot from the same cultures of the strain carrying the wt P_tet_-*fepA_-146+45_-lacZ* fusion used for the β-galactosidase assay. Strains used in this experiment are JJ389 and FQ99 (A), JJ389 and JJ330 (B, long fusion), and JJ135 and JJ146 (B, short fusion).

To go further, we next analyzed the same set of compensatory mutations in SdsR and *fepA* using the P_tet_-*fepA_-26+45_-lacZ* fusion (Fig. 3B). This simplified version of the *fepA* reporter is devoid of most of *fepA* 5’UTR and was previously shown to allow similar translational regulation to the full-length *fepA-lacZ* fusion, at least by sRNAs affecting *fepA* translation initiation, such as OmrA/B ((10) and see below). Repression of this shorter fusion by pSdsR was about 13-fold, which is reduced compared to the 36-fold repression of the fusion with the full-length 5’UTR. In addition, mutations SdsRmutβ or *fepAmutβ-lacZ* completely abolish repression, while combining these mutations to restore pairing also restored control (11-fold repression, Fig. 3B). This fully confirms that SdsR sRNA pairs to *fepA* TIR but also that full repression of *fepA* by SdsR requires an additional regulatory event that is dependent on a longer *fepA* 5’UTR. In summary, our data suggest that SdsR exerts a dual regulation on *fepA* expression, presumably through two distinct mechanisms.

### ArrS, RseX and SdsR are dependent on the long 5’UTR of *fepA* to control its expression

These results prompted us to dissect the role of the long *fepA* 5’UTR in the control by sRNAs in more detail. For this, we first determined its secondary structure using DMS and CMCT chemical probing, and an RNase A and T1 protection assay (Fig. S3). The reactivities of the different nts are summarized in Fig. 4A. The inferred structure is also supported by the cleavage pattern of *fepA* by lead-acetate, which is more prominent in all single-stranded regions (Fig. 4B). Briefly, several secondary structures are present in the *fepA* 5’UTR: a 5-base pair (bp) stem-loop at the very 5’ end (stem-loop I) and two imperfect stem-loops (II-b and II-c) forming a three-way junction with a 6-bp stem II-a (Fig. 4A). In contrast, the *fepA* TIR is very reactive to the different probes and thus appears mostly unstructured, consistent with our previous *in vitro* probing data (10).

**Figure 4.**
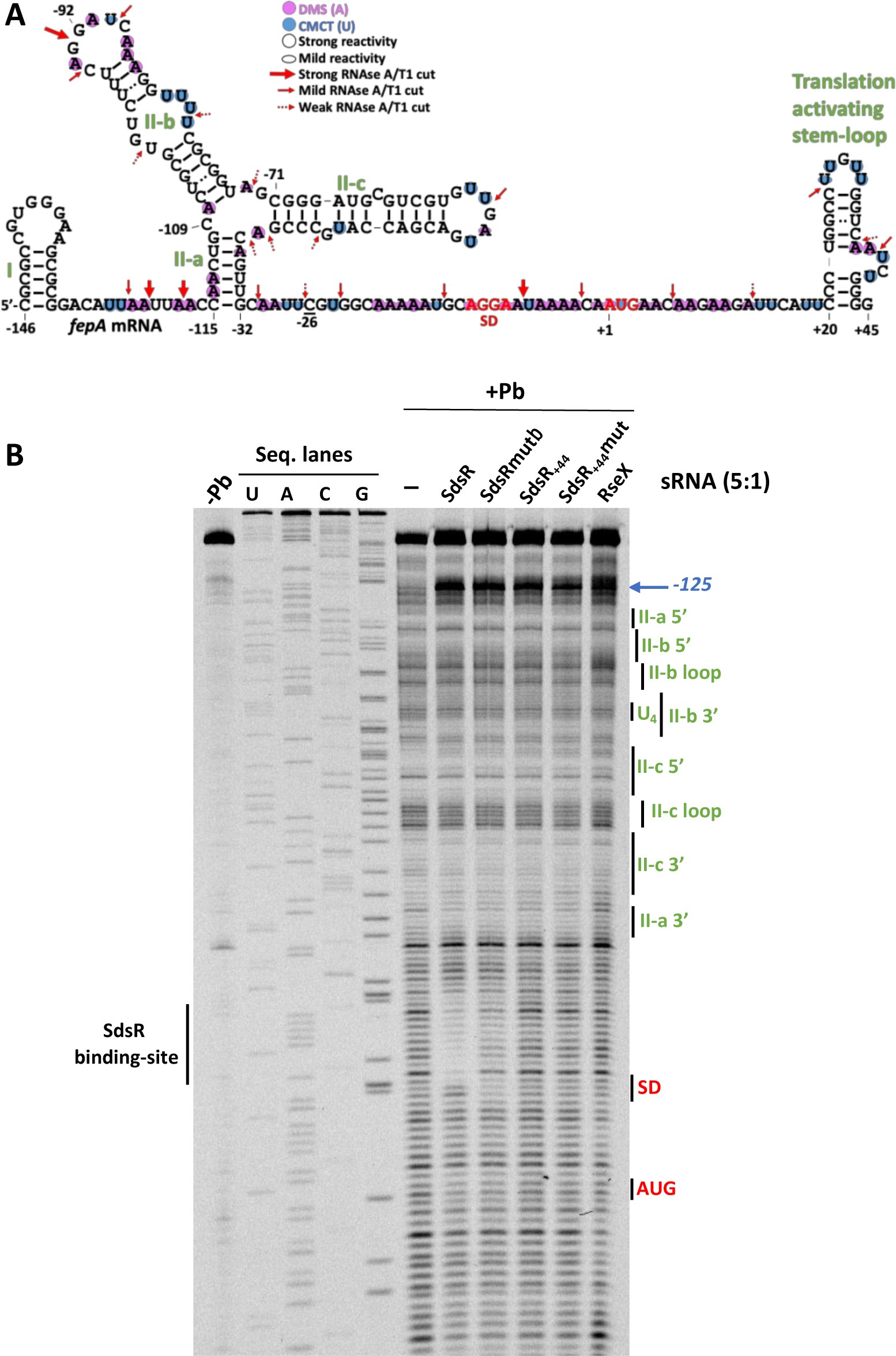
Secondary structure of the *fepA* 5’UTR *in vitro*. (A) Scheme of the structure of *fepA* 5’UTR based on the probing data with CMCT, DMS, and RNases A and T1 (Fig. S3), or with lead (panel B). The nomenclature of the different stem-loop structures is in green, and the SD and AUG start codon are in red. (B) Lead cleavage assay using an *in vitro* transcript corresponding to *fepA* mRNA region (-146+45). Cleavages were detected by reverse-transcription using the Cy5-labeled oligonucleotide FepAToeCy5_+26+45_ and the cDNA was analyzed on a sequencing gel, with the corresponding sequencing reactions (seq. lanes). When indicated, sRNAs were incubated with *fepA* mRNA at a 5:1 ratio prior to the lead treatment. -Pb: control lane without lead. The SdsR binding-site inferred from the protection from lead cleavage is shown on the left of the gel. Surprisingly, a very strong stop or pause in reverse-transcription was observed at position -125 relative to the start codon (blue arrow on the right of the gel) in the presence of the different sRNAs, regardless of their action on *fepA* expression *in vivo* and of their binding to *fepA in vitro*. We were not able to pinpoint what compound was responsible for this effect, but one possibility could be that *fepA* 5’UTR reacts to certain divalent ions or acetate, and that the sRNAs solutions interfere with this mechanism.

We next sought to assess the importance of the different structural elements in *fepA* 5’UTR for the control by sRNAs. To do so, we measured the repression of a set of *fepA-lacZ* fusions carrying either the full-length 5’UTR or truncated versions with only 120, 109, 92, 39 and 26-nt 5’UTR (Fig. 5A). These truncated constructs are expected to remove stem-loop I (construct 120), I and II-a (109), I, II-a and II-b (92), or all stem-loop structures in the 5’UTR (39 and 26). As expected, OmrA and OmrB sRNAs repressed *fepA* regardless of the length of its 5’UTR (Fig. 5A). The effects of the sRNAs RprA and RybB also appear largely independent of the long *fepA* 5’UTR. Indeed, RybB efficiently repressed all fusions, with a much higher repression factor for the (-26+45) version (>8-fold repression, compared to 2-to 4-fold for other fusions), while RprA similarly repressed fusions -146, -109, -92 and -39, but failed to repress the -120 and the -26 variants (Fig. 5A, left panel). This is easily explained for the -26 fusion, as this construct lacks part of the pairing region to RprA (Fig. 3A). In the case of the - 120 fusion, this could be due to a different accessibility of the RprA pairing region in this variant. The *fepA* 5’UTR is thus mostly dispensable for the regulation by OmrA, OmrB, RybB and RprA.

**Figure 5.**
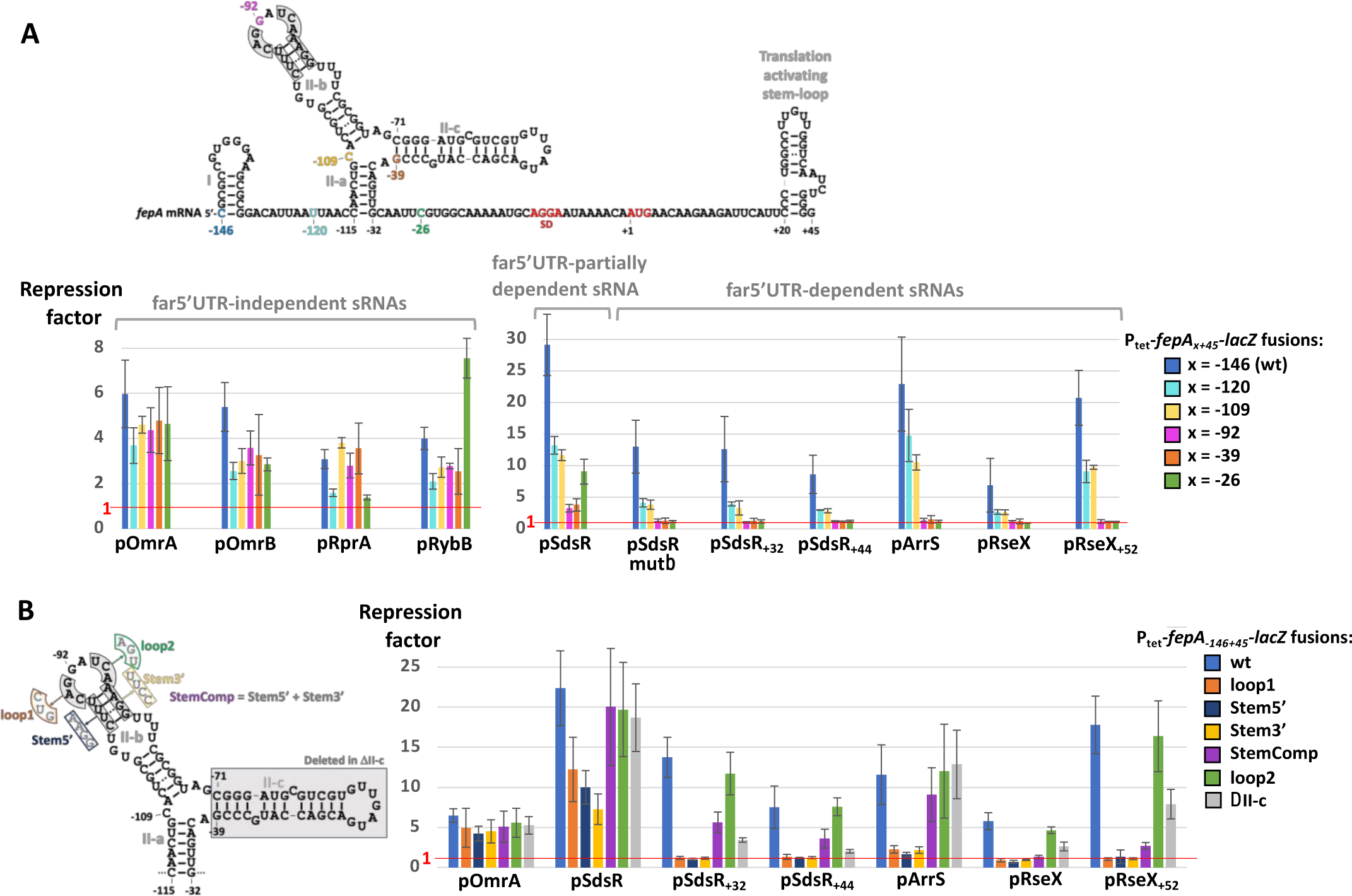
The role of *fepA* 5’UTR in the control by sRNAs. (A) The repression of *fepA* by the different sRNAs was analyzed for the derivatives of the P_tet_-*fepA-lacZ* fusion with shorter 5’UTR as indicated on the structure of the *fepA* (-146+45) region. For each sRNA, the repression factors were calculated relative to the vector control for the set of *fepA-lacZ* fusions carrying the full-length 5’ UTR (-146 fusion) or truncated versions with only the last 120, 109, 92, 39 or 26 nts. The red line indicates a repression factor of 1, i.e., regulation is completely abolished. Strains used in this experiment are JJ389 (-146), JJ406 (-120), JJ407 (-109), JJ136 (-92), FQ101 (-39) and JJ135 (-26). (B) As in (A) using mostly the far5’-UTR-dependent sRNAs, and mutants of the stem-loop II-b or II-c in the *fepA-lacZ* fusion as depicted on the structure scheme on the left of the graph. Strains used here are JJ389 (-146), JJ424 (loop1), MG2379 (Stem5’), MG2381 (Stem3’), MG2382 (StemComp), MG2385 (loop2) and MG2384 (ΔII-c).

The situation was completely different for the 3 other sRNAs repressing *fepA*. Indeed, control by ArrS, RseX or SdsR was reduced (roughly by half) for the –120 and -109 versions, and further reduced (for SdsR) or even abolished (for RseX or ArrS) when the 5’UTR was 92 nt-long or shorter (Fig. 5A, right panel). The demonstrated pairing of SdsR to the *fepA* TIR is likely responsible for the residual repression of the *fepA-lacZ* fusions -92, -39 or -26 by SdsR. We thus also measured repression of the full set of fusions by SdsRmutβ, i.e. the SdsR mutant that disrupts pairing to the *fepA* TIR (Fig. 3B). In this case, the pattern of regulation was highly similar to those observed with RseX and ArrS, as regulation was abolished when the 5’UTR of *fepA* was 92 nts or shorter (Fig. 5A). This confirms that SdsR represses *fepA* both via pairing to the mRNA TIR and via an independent regulatory process that requires the upstream portion of the *fepA* 5’UTR. For simplicity, we will distinguish between these two control mechanisms as far5’UTR-independent and far5’UTR-dependent, respectively, for the rest of the manuscript.

Our results clearly show a role for the promoter-proximal portion of the *fepA* 5’UTR in the repression by SdsR, RseX and ArrS, and we therefore wondered whether it was sufficient for these control processes or not. To address this question, we used another set of *fepA*-lacZ fusions, truncated this time from the 3’ end of the *fepA* moiety and where *lacZ* is fused in frame after nt +30 or +3 of *fepA*, or fused after nt -26 and carrying the *lacZ* TIR in this last case (Fig. S4, panels A and B). None of the far5’UTR-dependent or -independent sRNAs was able to significantly control the (-146-26) fusion, clearly showing that the *fepA* TIR is required for all these control mechanisms (Fig. S4B). Moreover, the fusion ending at *fepA* nt +3 was controlled to varying degrees by the far5’UTR-dependent sRNAs, showing that the beginning of the coding sequence plays a role as well, albeit reduced. Finally, the fusion with *fepA* ending at +30 was still controlled by the different sRNAs, except for OmrA and OmrB, as expected since this construct removes the 3’ side of the translation activating structure that is the target of these sRNAs (10).

Together, these data show that, of the seven sRNAs that repress *fepA*, the three that had the strongest effect when overproduced are dependent on the upstream portion of the *fepA* 5’UTR. At this stage however, their mode of action on *fepA* was still unclear and we focused on two of them, RseX and SdsR, to address this question.

### Repression of *fepA* by the 3’-ends of SdsR or RseX sRNAs

SdsR is transcribed as a 104 nts-long sRNA and can be processed by RNase E, producing a shorter sRNA devoid of the first 31 nts (39). This processed form is hereafter referred to as SdsR_+32_. SdsR was shown to target multiple mRNAs, including *ompD* in *Salmonella* (39), *mutS* (40), *tolC* (41) and two pairing regions have been identified within this sRNA (42). Interestingly, the first pairing region of SdsR is the one that interacts with the *fepA* TIR, and it is absent in the processed SdsR_+32_ (Fig. 6A). As data from Fig. 3 and Fig. 5 show a repressive effect of SdsR on *fepA* independently of its pairing to the TIR, we asked whether SdsR_+32_ could repress *fepA*. Using the P_tet_-*fepA_-146+45_-lacZ* fusion as a reporter, we found that SdsR_+32_ down-regulated *fepA* 16-fold. As expected, this is lower than the repression by full-length SdsR (22-fold, Fig. 6B), as the pairing to the TIR is lost with SdsR_+32_. To test the involvement of the second pairing region of SdsR in *fepA* regulation, we constructed a second truncated version of SdsR, namely SdsR_+44_, that lacks both previously identified pairing regions (Fig. 6A) and found that it still repressed *fepA*, by a factor of 9-fold. Finally, we also tested these two short forms of SdsR on the set of *fepA-lacZ* reporters with different 5’UTR lengths. Their regulation pattern was strikingly similar to that of SdsRmutβ, with repression of fusions with shorter 5’UTR being impaired or abolished (Fig. 5A). These data demonstrate that the last 63 nts of SdsR are necessary and sufficient for the regulation of *fepA* that is independent of the TIR-pairing.

**Figure 6.**
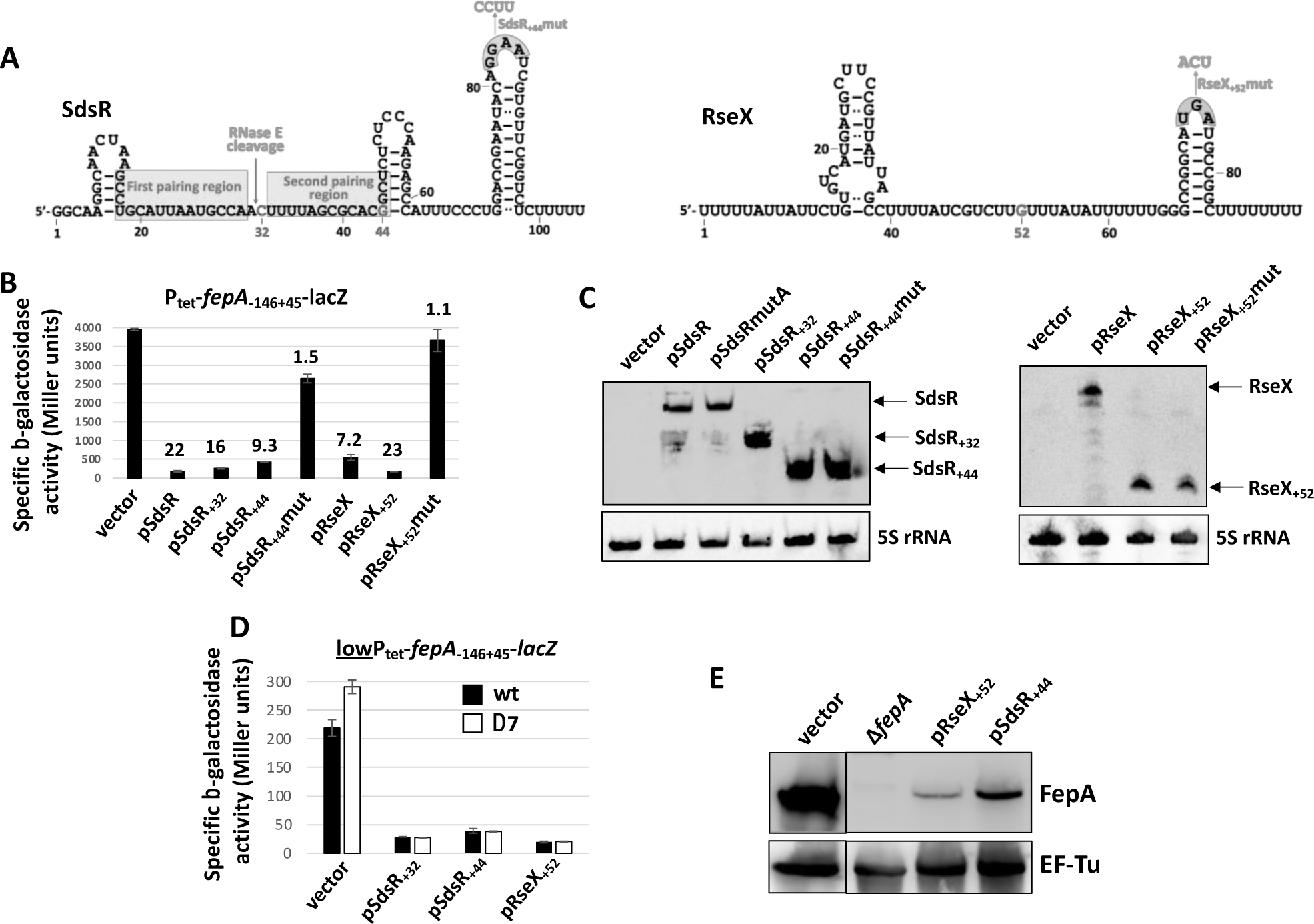
The terminator regions of RseX and SdsR repress *fepA*. (A) Secondary structure of SdsR and RseX, as depicted in (42) and (64), respectively. The position of the RNase E cleavage in SdsR and of the two pairing regions is also from (42). The nts in red indicate the 5’ ends of the short forms of SdsR or RseX used in this study, while the mutations introduced in the terminator loops are shown in grey. (B) The β-galactosidase activity of the P_tet_-*fepA_-146+45_-lacZ* fusion (strain JJ389) was measured in the presence of these different SdsR or RseX derivatives, and (C) the levels of these sRNAs were analyzed by Northern blot from the same cultures. To simultaneously detect RseX and its variants, the probes complementary to wt RseX or RseX_+52_mut were mixed at a 1:1 ratio. (D) Effect of the short SdsR and RseX variants on the expression of the lowP_tet_-*fepA_-146+45_-lacZ* fusion in the wt (strain OK410) or the Δ7 background (strain ES250). (E) Western blot analysis of FepA levels in the *Δfur* strain ES021 grown to exponential phase with the indicated plasmids. The EF-Tu protein was used as a loading control in this experiment. For clarity, only relevant lanes of the blot are shown, with the black line indicating the junction between two regions of the same membrane. The full membrane is presented in Fig. S5.

Since control of *fepA* by RseX and ArrS also displayed a profile similar to that of SdsRmutβ, SdsR_+32_ and SdsR_+44_ (Fig. 5A), we wondered whether the 3’-end of these sRNAs could be responsible for *fepA* regulation as well. This was tested for RseX, by using a short form of this sRNA devoid of the first 51 nts, hereafter named RseX_+52_. Strikingly, this variant was even more efficient than full-length RseX in controlling *fepA*, as it repressed *fepA-lacZ* 23-fold, compared to 7-fold for wt RseX. Furthermore, similar to the other far5’UTR-dependent sRNAs (SdsRmutβ, SdsR_+32_, SdsR_+44_, ArrS and RseX), reducing the 5’UTR to 92 nts or less completely abolished control by RseX_+52_ (Fig. 5A). Thus, the 3’-ends of both SdsR and RseX are responsible for the far5’UTR-dependent control of *fepA*. This unexpected result raises several intriguing questions, in particular which common feature(s), if any, of SdsR(_+44_) and RseX(_+52_) may mediate *fepA* control, and by which mechanism.

### SdsR and RseX transcriptional terminator loops drive *fepA* regulation, likely without base-pairing

The most obvious common element between SdsR_+44_ and RseX_+52_ is the presence of a Rho-independent transcriptional terminator, and we thus wondered whether they could mediate *fepA* control. As all sRNAs tested in our original screen (Fig. 1A), including the ones that did not regulate *fepA* expression, also possess such terminators, it is highly unlikely that the stem-loop structure or the stretch of Us could explain the effect of SdsR_+44_ and RseX_+52_. For this reason, we focused on the exposed nts of the terminator loop and tested their importance in *fepA* control by mutating some of them in SdsR_+44_ (mutant SdsR_+44_mut) and RseX_+52_ (mutant RseX_+52_mut). Strikingly, these two mutant sRNAs strongly impaired *fepA* regulation, even though they accumulated to a comparable level to the un-mutated SdsR_+44_ and RseX_+52_ (Fig. 6B). This indicates that, at least for these two sRNAs, the loops of the intrinsic terminators are involved in gene regulation, which is rather unusual.

We were not able to predict a convincing base-pairing interaction between SdsR_+44_ or RseX_+52_ and the *fepA* mRNA. Nonetheless, we could not rule out such a pairing on this basis alone. To test for a possible direct interaction, we analyzed the lead cleavage pattern of an *in vitro* transcribed *fepA* mRNA fragment encompassing the 5’ UTR and the early coding sequence, in the presence of several sRNAs: wt SdsR, SdsRmutβ, SdsR_+44_, SdsR_+44_mut and wt RseX. While the addition of wt SdsR protected the (-22-11) region of *fepA* mRNA, in complete agreement with the pairing shown in Fig. 3B, none of the other sRNAs tested modified the lead cleavage pattern of *fepA* (Fig. 4B). These data suggest that repression of *fepA* by RseX or SdsR is likely to involve additional factor(s) and may not rely only on a classical base-pairing mechanism between *fepA* mRNA and these sRNAs.

### Stem-loop II-b of the *fepA* 5’UTR is required for the control by SdsR, RseX and ArrS sRNAs

We next wondered which feature of the *fepA* mRNA is involved in the control by the far5’UTR-dependent sRNAs. As these sRNAs can still repress the -109 *fepA-lacZ* fusion, but not the -92 or shorter fusions (Fig. 5A), we reasoned that the stem-loop II-b most likely plays an important role. This was tested more directly by introducing mutations in the exposed nts -95 to -88 of the apical loop (mutations Loop1 and Loop2) or in the upper part of the stem (mutations Stem5’ and Stem3’ to open this structure, and the compensatory mutation StemComp) (Fig. 5B). Strikingly, the Loop1, Stem5’ and Stem3’ mutations completely abolished the regulation of *fepA* by all the far5’UTR-dependent sRNAs, confirming the key role of stem-loop II-b. Furthermore, the restoration of the upper stem structure with the StemComp variant fully restored control by ArrS, and partially by SdsR_+32_ or SdsR_+44_, showing the importance of the secondary structure rather than the sequence of the stem for these sRNAs (Fig. 5B). In contrast, RseX or RseX_+52_ barely affected the expression of the StemComp fusion, suggesting there may be contextual differences between the modes of action of RseX, SdsR and ArrS. Lastly, the Loop2 change had no noticeable effect at all, contrasting with the major effect of the Loop1 mutation. This suggests that only specific nts of the apical loop II-b are important for sRNA control. In full agreement with the previous data, the control by SdsR followed the same pattern as SdsR_+32_ or SdsR_+44_, except that the regulation was reduced, and not abolished, in the Loop1, Stem5’ or Stem3’ mutants, consistent with the fact that the pairing of SdsR to the TIR is not affected in these 5’UTR mutants. Similarly, we found that the far5’UTR-independent sRNA OmrA repressed the wt and the different mutants of *fepA* equally efficiently (Fig. 5B).

We also tested the importance of other elements of *fepA* mRNA, and in particular the stem-loop II-c, by deleting it in the *fepA-lacZ* reporter fusion (ΔII-c construct, Fig. 5B), as well as the sequence around the SD and AUG by using the mutβ and mutγ variants (Fig. S4A). These changes impaired control by some of the different far5’UTR-dependent sRNAs, such as SdsR_+32_, SdsR_+44_, RseX, RseX_+52_ or ArrS, but never as drastically as changes in the stem-loop II-b (Fig. 5B and S4C).

In conclusion, stem-loop II-b is a major determinant of the *fepA* mRNA for the control by the ArrS, RseX and SdsR sRNAs that are likely to act on the *fepA* 5’UTR independently of a direct base-pairing interaction. Apart from minor differences between them (for instance, with the StemComp variant), the similar regulation patterns of the different stem-loop II-b mutants suggests that a common factor likely mediates their effects on *fepA*.

### Hfq and RNase E are required for *fepA* control by ArrS, RseX and SdsR

RNA-binding proteins (RBPs) are obvious candidates for factors involved in gene control by sRNAs, and several of them, among which Hfq, ProQ, CsrA or different ribonucleases were previously found to be essential for sRNA action (*e.g.* 31, 32, 42–44). Hence, we tested whether the different sRNAs still control *fepA* in strains mutated for some RBPs. While the control of *fepA* was not affected in mutants of the genes for ProQ, CsrA, RNase III or PNPase (data not shown), we found that Hfq and the RNase E were in contrast involved in these controls.

To study the role of Hfq, we used not only a deletion of the *hfq* gene, but also mutants of three different RNA binding surfaces of the Hfq protein that have been described previously: the proximal face (Q8A mutant), the distal face (Y25D) or the rim (R16A) (46–48). The proximal face recognizes U-rich sequences, typically found in intrinsic terminators, and interacts with a predominant class of Hfq-binding sRNAs (also referred as Class I sRNAs), while the distal face recognizes successive (AAN) motifs, often found in mRNAs that are targeted by Class I sRNAs. The lateral rim surface makes additional contact with different RNA substrates via a patch of arginine residues. Because the deletion of *hfq* or the point mutations in Hfq proximal face (Q8A) or rim (R16A) drastically decreased the levels of most of the sRNAs used in this study (Fig. S6A), we mostly focused on the distal face mutant Y25D. In these experiments, we assessed the expression of *fepA* using the P_tet_-*fepA*-*mSc* fusion and we found that only the full-length SdsR repressed *fepA* in the *hfq*Y25D mutant (Fig. 7A). This is consistent with the *in vitro* observation that the pairing of SdsR to the SD region of *fepA* exists in the absence of any other factor (Fig. 4B). In contrast, repression by all other sRNAs, including SdsR_+44_, was abolished in the *hfq*Y25D context, despite the fact that all sRNAs accumulate to higher levels in this mutant background compared to the wt strain (Fig. 7B).

**Figure 7.**
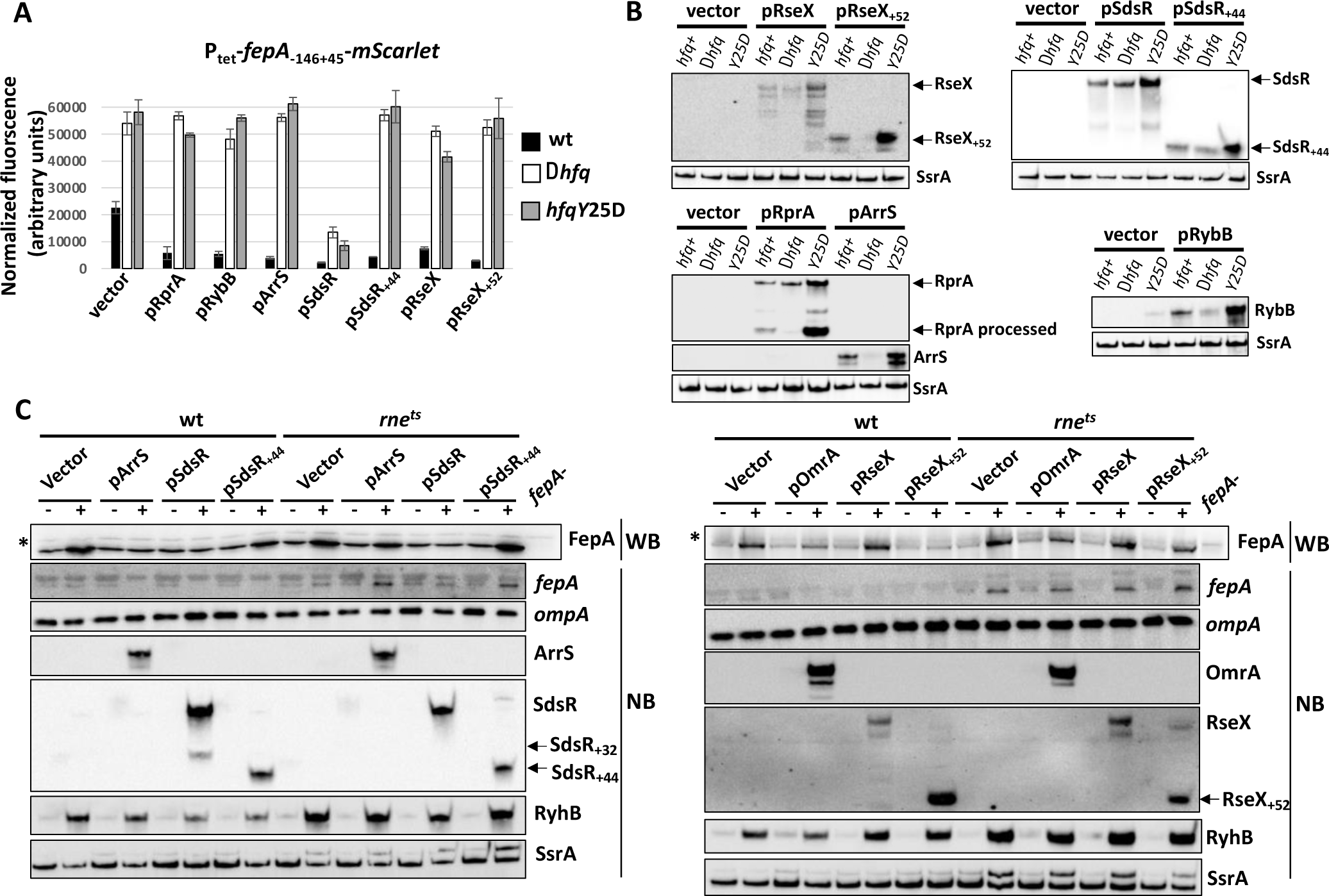
Control of *fepA* by the 5’ UTR-dependent sRNAs depends on Hfq and RNase E. (A) The *fepA-mSc* fluorescence was measured in the *hfq+*, (MG2412), *Δhfq* (MG2413) and *hfqY25D* (MG2416) strains upon overexpression of RprA, RybB, ArrS, SdsR, SdsR_+44_, RseX and RseX_+52_ sRNAs. OmrA and OmrB were omitted from this analysis since they do not significantly repress this reporter fusion. (B) The levels of the overproduced sRNAs were analyzed by Northern blot from cultures in CAG-tetracycline-IPTG 250 μM, i.e., the same medium as used for the fluorescence measurement in (A). (C) The control of *fepA* by ArrS, SdsR and SdsR_+44_ (left panel) or by OmrA, RseX and RseX_+52_ (right panel) was analyzed in a wt or an *rne^ts^* background. For this, cells were grown at 30°C to exponential phase and an aliquot was shifted to the non-permissive temperature of 44°C for 15 minutes in the presence of 250 μM 2, 2’-dipyridyl and 100 μM IPTG to induce *fepA* and the sRNA transcription, respectively (+ lanes). As a control, growth of cells continued at 30°C for 15 minutes, without 2, 2’-dipyridyl or IPTG (-lanes). RNA and proteins were then extracted and Northern or Western blot analysis of specific candidates was performed. Strains used in this experiment are MG1325 (*rne^+^*) and MG1326 (*rne^ts^*). The asterisk indicates a non-specific band detected with the FepA antibody, and used here as a loading control.

Expression of *fepA* was strikingly up-regulated in the *Δhfq* or *hfqY25D* mutant (Fig. 2B; Fig. 7A, vector lanes), and one possibility to explain the effect of the 3’ ends of RseX or SdsR is that they would generally enhance Hfq action on mRNA-targets. Since the expression of the *mutS* mRNA is also strongly up-regulated (about 10-fold) in either *Δhfq* or *hfqY25D* strains (49), we analyzed the impact of ArrS, RseX, SdsR and their short forms on a *mutS-lacZ* fusion. Consistent with previous reports (49), this fusion was repressed 3-fold by ArcZ, used here as a positive control, and, to a lesser extent, by SdsR (1.8-fold repression) (Fig. S6B). However, we find that its expression is barely modulated by SdsR_+44_ (1.2-fold repression) and completely insensitive to ArrS, RseX or RseX_+52_ overproduction. Thus, these sRNAs do not have a general effect on the expression of Hfq-regulated genes.

We also studied a possible involvement of the essential RNase E endonuclease in *fepA* control. For this, we inactivated a thermosensitive RNase E enzyme encoded by the *rne^ts^* allele by shifting cells to non-permissive temperature for 15 minutes, while transcription of endogenous *fepA* mRNA and of the different plasmid-borne sRNAs was induced by addition of 2-2’-dipyridyl and IPTG, respectively (Fig. 7C, + lanes). In parallel, the same experiment was also performed with wt cells, and a negative control where cells were kept at permissive temperature without *fepA* or sRNA induction was done systematically (-lanes). Proteins and RNAs were then extracted and analyzed by Western or Northern blots, respectively. With this experimental set-up, we observed clear regulation of FepA levels by ArrS, SdsR, SdsR_+44_ and RseX_+52_ in wt cells. In the *rne^ts^*context, regulation by ArrS was reduced and regulation by RseX_+52_ and SdsR_+44_ was abolished. In contrast, SdsR still decreased FepA protein levels in the *rne^ts^* mutant, even though *fepA* mRNA levels were not affected by SdsR in this background. Of note, more modest regulations of *fepA*, for instance by OmrA or wt RseX, were not easily assessed in this experiment, even in the wt strain, and deciphering the role of RNase E in these cases would require to adjust the experimental conditions (see for instance (10) for OmrA). In any case, these data clearly point to a role for both Hfq and the RNase E in the control of *fepA* by the far5’UTR-dependent sRNAs.

## DISCUSSION

This work reveals that at least five sRNAs, in addition to the previously studied OmrA and OmrB, repress the expression of the *fepA* gene in *E. coli*. Even more sRNAs could be involved in *fepA* control, either because they were not present in the plasmid library used in this study, or because, as seen for OmrA and OmrB sRNAs, the *fepA-mSc* reporter that was used here to identify them is not a suitable read-out for their action (Fig. 1). Except for OmrA and OmrB which are at least partially redundant, these seven sRNAs respond to different environmental cues via different transcriptional regulators (Fig. 8 and references therein). They can nonetheless be co-expressed, for instance when cells reach stationary phase (Fig. 2C). These results thus suggest that it is beneficial for the cell to limit FepA levels under different conditions. Intuitively, these regulatory mechanisms could be more efficient when *fepA* transcription is less abundant, such as under iron-replete conditions, and they could participate to limit iron toxicity. Alternatively, these sRNAs could limit *fepA* de-repression by Fur under conditions combining iron starvation and FepA toxicity, such as those induced by phage H8 (50) or colicins B or D (51). Further investigation could reveal whether any of these sRNAs itself respond to iron, phages and/or colicins, and how these regulations articulate with the Fur-mediated transcriptional control of *fepA*. It would also be interesting to determine if this repression of *fepA* by multiple sRNAs participates to the different processes that are controlled by changes in iron levels, among which antibiotics resistance (52, 53) or the iron-based memory impacting swarming motility (54).

**Figure 8.**
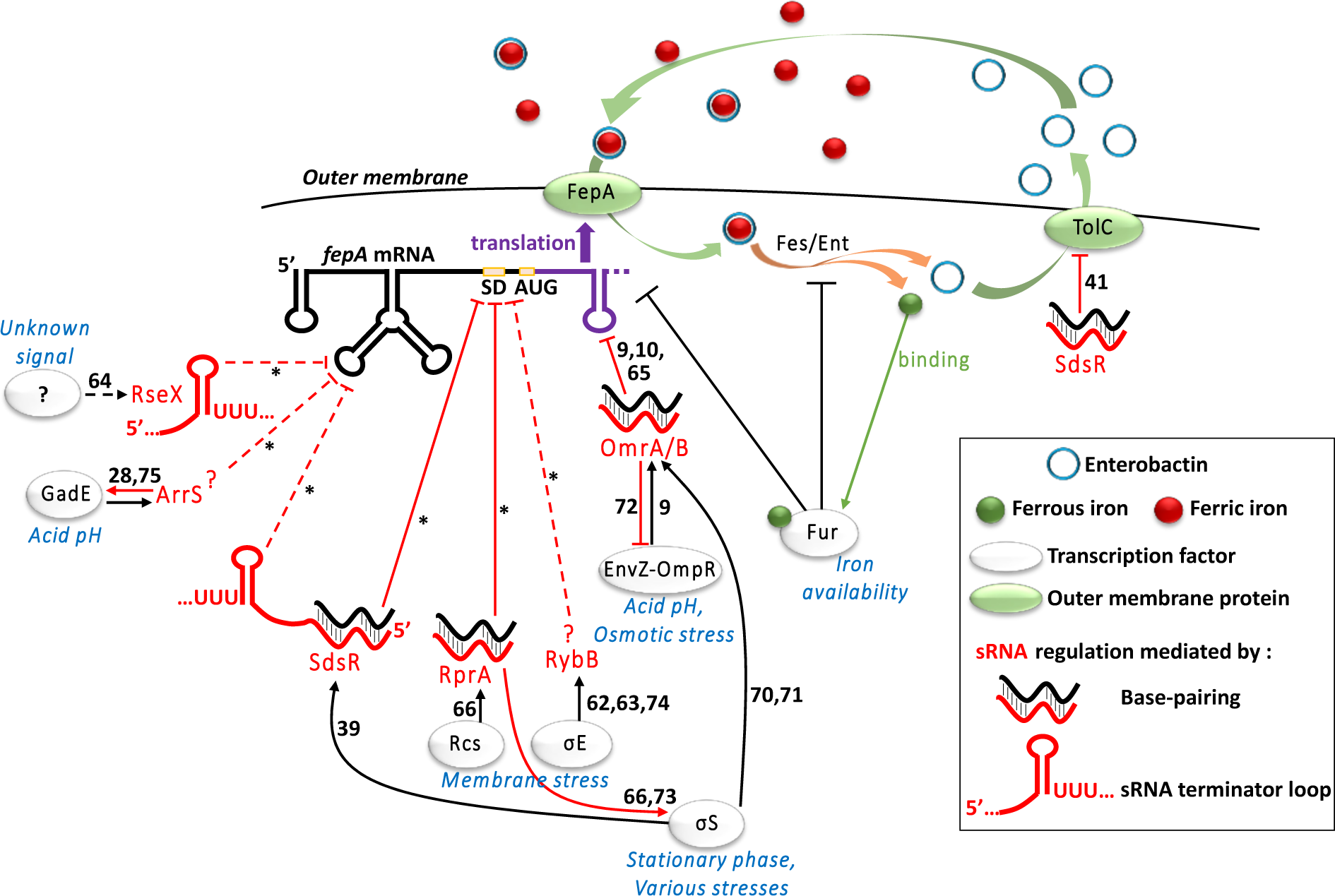
Model of the regulation of *fepA* by multiple sRNAs, in response to different signals. Transcriptional regulators and transcriptional controls are shown in black, and sRNAs and the post-transcriptional controls are in red, with the dotted lines referring to possibly indirect regulations and/or that involve additional factor(s). The physiological signals and environmental cues impacting the activity of the different transcription factors are in blue. The numbers in black provide the references for the previously described regulations, while the asterisks indicate regulations unraveled in this study.

Another key finding made in this study is that, in some cases, the intrinsic transcriptional terminators of the sRNAs are required for their regulatory action. These terminators were already known to be strong Hfq binding sites via their long U stretches (55, 56), but we report here that the sequence of the exposed nts can have an important role as well (Fig. 6). This is reminiscent of what was shown for the OxyS sRNA in the control of the gene encoding the FhlA transcriptional activator, but with the major difference that the terminator loops of the sRNAs apparently act via different mechanisms to control *fhlA* or *fepA*. Indeed, the exposed nts of the OxyS terminator loop are directly involved in the pairing with the mRNA-target as they form a kissing complex with the *fhlA* TIR, while a second kissing loop interaction is made between another loop of OxyS and the *fhlA* coding sequence (57, 58). In contrast, our results do not support a direct interaction of SdsR or RseX terminator loops with the *fepA* mRNA (Fig. 4), but they could pair instead to another sRNA or mRNA, whose regulation would in turn affect *fepA* expression.

Alternatively, they could be involved in the interaction with other factors, including Hfq itself. However, because the mutants of the terminator loop (RseX_+52_mut and SdsR_+44_mut), accumulate to levels comparable with their un-mutated counterparts (RseX_+52_ and SdsR_+44_), it is unlikely that they are strongly affected in their ability to bind Hfq. Furthermore, the effect of the far5’UTR-dependent sRNAs does not appear to be simply due to a general effect on Hfq as the known Hfq target *mutS* does not respond to these sRNAs (Fig. S6B).

In addition to its RNA binding activities, Hfq is involved in numerous protein-protein interactions, many of which are RNA-dependent ((59) and references therein). Hence, one can hypothesize that mutating the exposed nts of RseX_+52_ or SdsR_+44_ changes the ability of Hfq to interact with its other partners. Of note, RNase E is one of the proteins that interacts with Hfq in an RNA-dependent manner (59), and it would be interesting to determine if RseX_+52_ or SdsR_+44_ somehow affects the formation of Hfq-RNase E complexes. More generally, understanding how Hfq and RNase E, as well as other potential factors, are involved in the sRNA control of *fepA* will be useful to explain the specificity of these regulatory mechanisms for which no sRNA-mRNA pairing could be demonstrated. This situation is indeed different from the other examples of RNase E-dependent control by sRNAs reported previously for which target specificity was provided by sRNA-mRNA pairing (43, 60, 61). At this stage, we cannot exclude that some of the effects that we observe on *fepA* are due to an overproduction of the sRNAs. Even if it were the case, understanding the underlying mechanism is likely to bring important insight into the post-transcriptional regulation of this important gene for iron acquisition.

Being the receptor of iron-siderophore complexes, FepA thus belongs to two classes of proteins whose synthesis has been repeatedly found to be subjected to sRNA control: the outer membrane proteins and the proteins involved in iron homeostasis. Some of the sRNAs involved in these controls of outer membrane proteins or iron-related genes were notably RybB (62, 63), SdsR (39, 41, 42), RseX (64), OmrA or OmrB (9, 10, 65), and the findings that they also repress *fepA* expression seems thus consistent regarding their other targets. The *fepA* mRNA furthermore possesses a long 5’UTR, which is also the case of other mRNAs controlled by multiple sRNAs such as *csgD*, *flhDC* or the *rpoS* mRNA, whose 5’UTRs are 148-, 198- and 567-nt long, respectively. Again, some of the *fepA* repressors, namely OmrA, OmrB, RybB and RprA, are also regulators of *csgD* (14–16), *flhDC* (18) or *rpoS* (66). The interaction between Hfq and the exceptionally long *rpoS* 5’UTR was analyzed in detail and Hfq was found to bind both an AAN repeated motif and a U-rich loop of the *rpoS* leader, resulting in a compact structure of the mRNA when in complex with Hfq and the exposure of the sRNA binding-site (67). Interestingly, *fepA* 5’UTR also presents both a U-rich loop in the vicinity of stem-loop II-b and successive (AAN) motifs in the TIR, *i.e.*, in two regions that are required for control by the far5’UTR-dependent sRNAs (Fig. 5 and S4). Hence, an exciting hypothesis is that Hfq could similarly contact these two sites of the *fepA* leader and, if true, understanding the outcomes for the sRNA regulation will be of great interest. Furthermore, studying in detail the interaction between Hfq and the RseX sRNA also seems appealing given the presence of U stretches on both the 5’ and 3’ sides of the RseX transcriptional terminator (Fig. 6B), which could both play a role in contacting Hfq.

The *csgD*, *flhDC*, *rpoS* and *fepA* mRNAs thus provide examples of long 5’UTRs allowing control by multiple sRNAs. While there are numerous examples of mRNAs with shorter 5’UTRs targeted by sRNAs at their TIR and/or within their coding sequence downstream of the TIR, *e.g.* the OmpD porin gene in *Salmonella* (39, 61, 62, 68, 69), *fepA*, *csgD* and *flhDC* are so far the mRNAs with the most numerous sRNA regulations described in bacteria. In particular, *fepA* 5’UTR provides a functional and efficient hub for post-transcriptional regulation, in addition to the *fepA* TIR and coding region. Overall, this raises the question of whether mRNAs with long 5’UTRs are more represented among sRNA-regulated genes, or among genes with specific functions that require multiple regulatory pathways, such as membrane proteins genes for instance. More generally, whether a correlation exists between the length of mRNA 5’UTRs and the number of post-transcriptional regulators remains to be determined.

## ACKNOWLEDGMENTS

We thank P. Mandin and S. Gottesman for the original sRNA plasmid library, as well as N. Majdalani and J. Chen for several *E. coli* strains, and K. Postle for the anti-FepA antibody. We are grateful to members of the group for discussions during this project and to C. Condon for critical reading of the manuscript.

