## Supplementary data for "Control of iron acquisition by multiple small RNAs unravels a new role for transcriptional terminator loops in gene regulation"

A

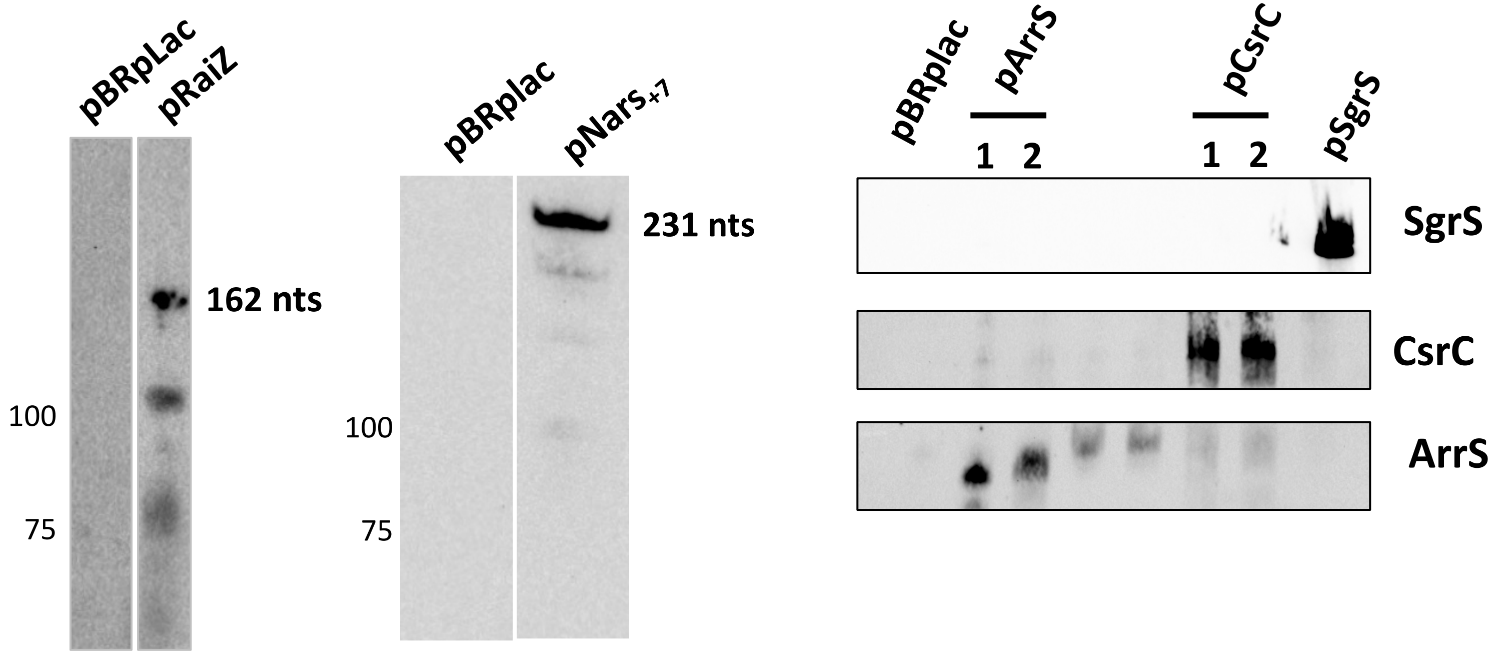

**Fig. S1. The sRNA overproduction was verifed by Northen blot for the plasmids constructed in the course of this study.** RNA was extracted from the cloning strain NEB5alpha transformed with the indicated plasmids and grown to exponential phase in LB-Amp-IPTG medium. Only the relevant lanes of the blots are annotated.

A

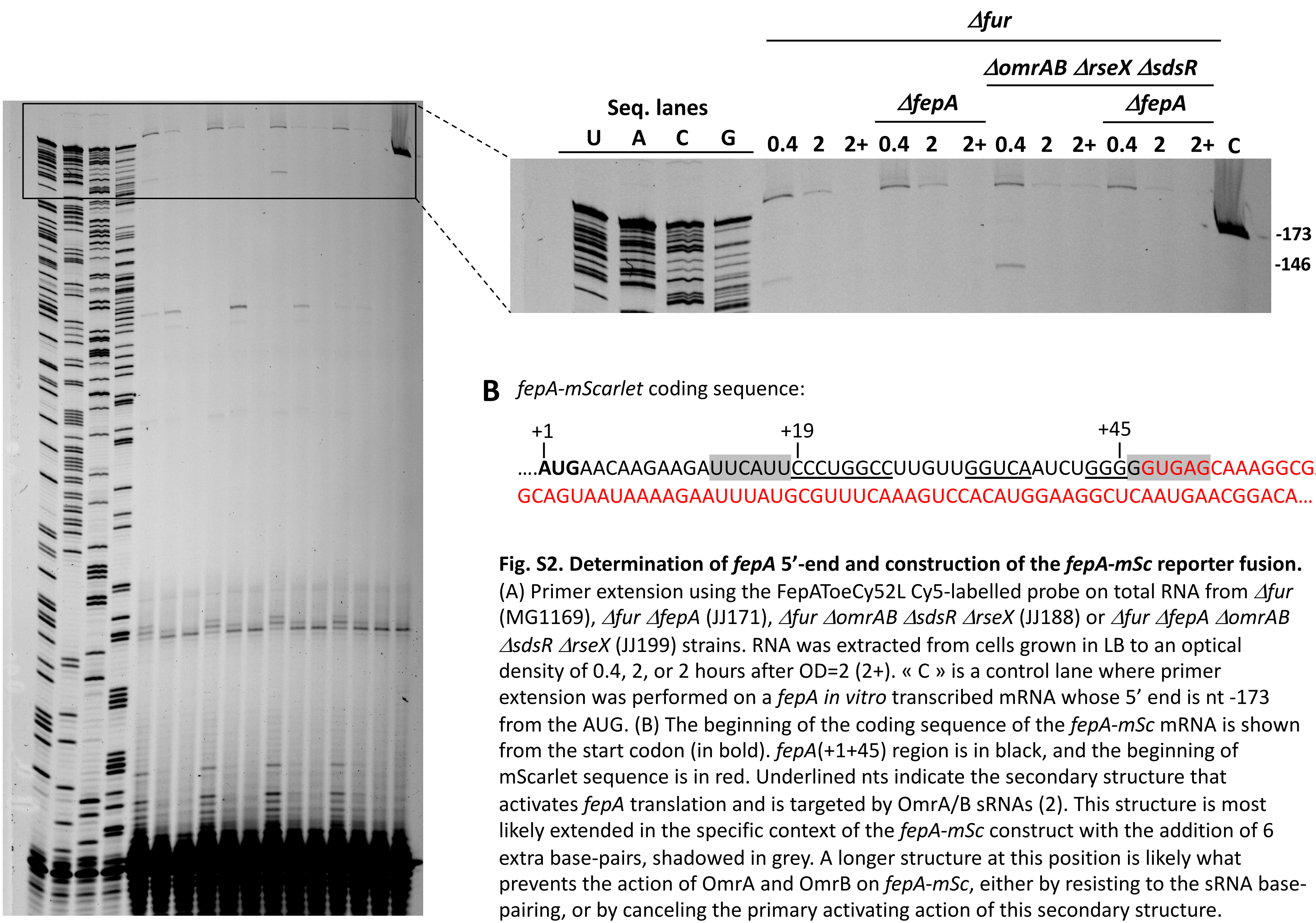

**B** *fepA-mScarlet* coding sequence:

+1 +19 +45

....**AUG**AACAAGAAGAUUCAUUCCCUGGCCUUGUUGGUCAAUCUGGGGGUGAGCAAAGGCGAA
GCAGUAAUAAAAGAAUUUAUGCGUUUCAAAGUCCACAUGGAAGGCUCAAUGAACGGACA...

**Fig. S2. Determination of *fepA* 5'-end and construction of the *fepA-mSc* reporter fusion.** (A) Primer extension using the FepAToeCy52L Cy5-labelled probe on total RNA from *Δfur* (MG1169), *Δfur ΔfepA* (JJ171), *Δfur ΔomrAB ΔsdsR ΔrseX* (JJ188) or *Δfur ΔfepA ΔomrAB ΔsdsR ΔrseX* (JJ199) strains. RNA was extracted from cells grown in LB to an optical density of 0.4, 2, or 2 hours after OD=2 (2+). « C » is a control lane where primer extension was performed on a *fepA in vitro* transcribed mRNA whose 5' end is nt -173 from the AUG. (B) The beginning of the coding sequence of the *fepA-mSc* mRNA is shown from the start codon (in bold). *fepA*(+1+45) region is in black, and the beginning of mScarlet sequence is in red. Underlined nts indicate the secondary structure that activates *fepA* translation and is targeted by OmrA/B sRNAs (2). This structure is most likely extended in the specific context of the *fepA-mSc* construct with the addition of 6 extra base-pairs, shadowed in grey. A longer structure at this position is likely what prevents the action of OmrA and OmrB on *fepA-mSc*, either by resisting to the sRNA base-pairing, or by canceling the primary activating action of this secondary structure.

Fig. S2

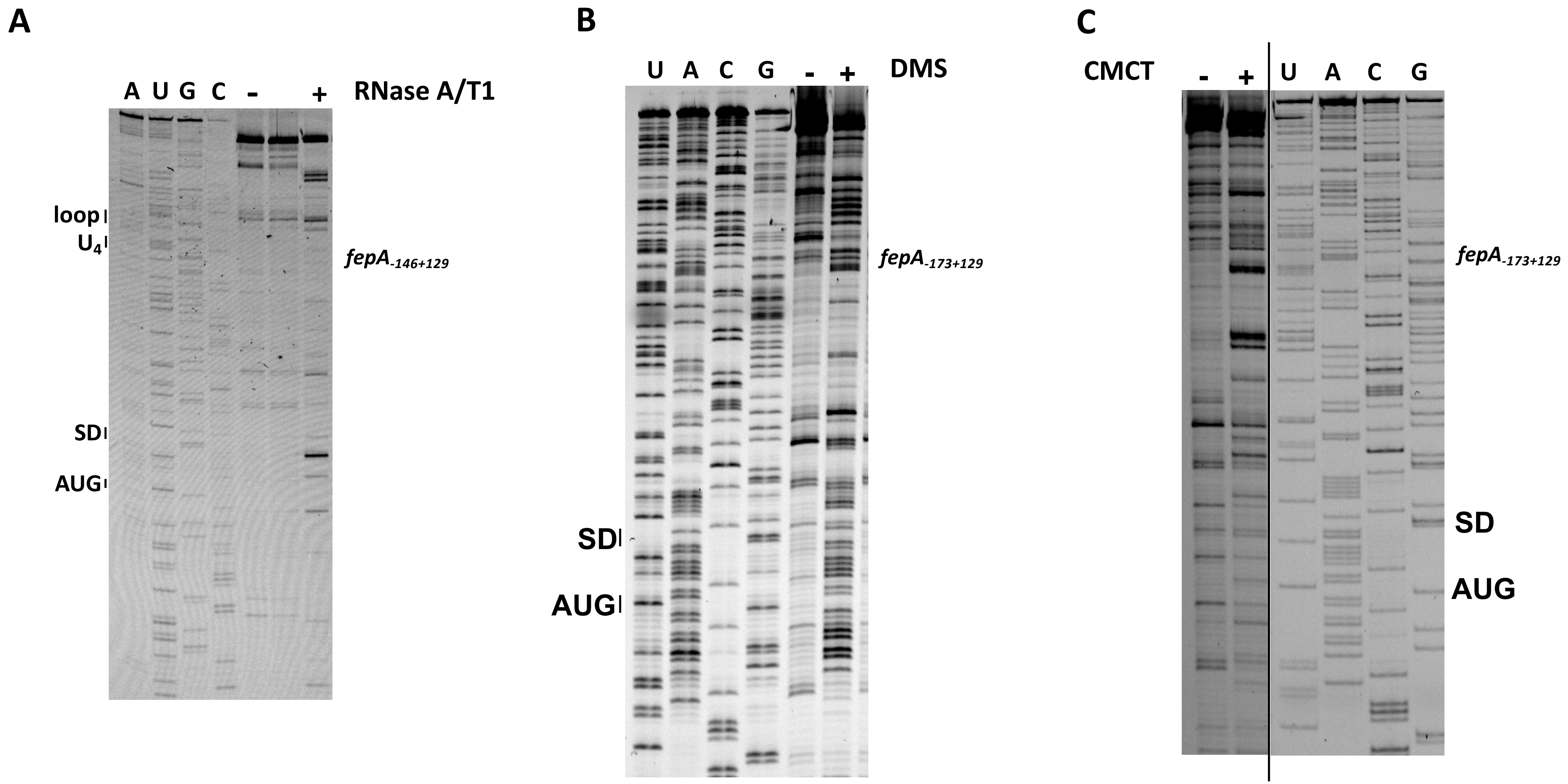

**Fig. S3. Determination of *fepA* mRNA secondary structure.** (A) *fepA in vitro* transcript corresponding to the (-146+129) region of the mRNA was incubated with a mix of RNases A and T1 before primer extension with the Cy-5 labelled primer FepAToeCy5 and separation on a sequencing gel. Only the relevant lanes are annotated. (B, C) The *fepA* in vitro transcript (-173+129) was subjected to modification by DMS (methylates A and C residues, panel B) or CMCT (modifies U, and to a lesser extent, G residues, panel C); modifications were then analyzed by primer extension as in (A).

Fig. S3

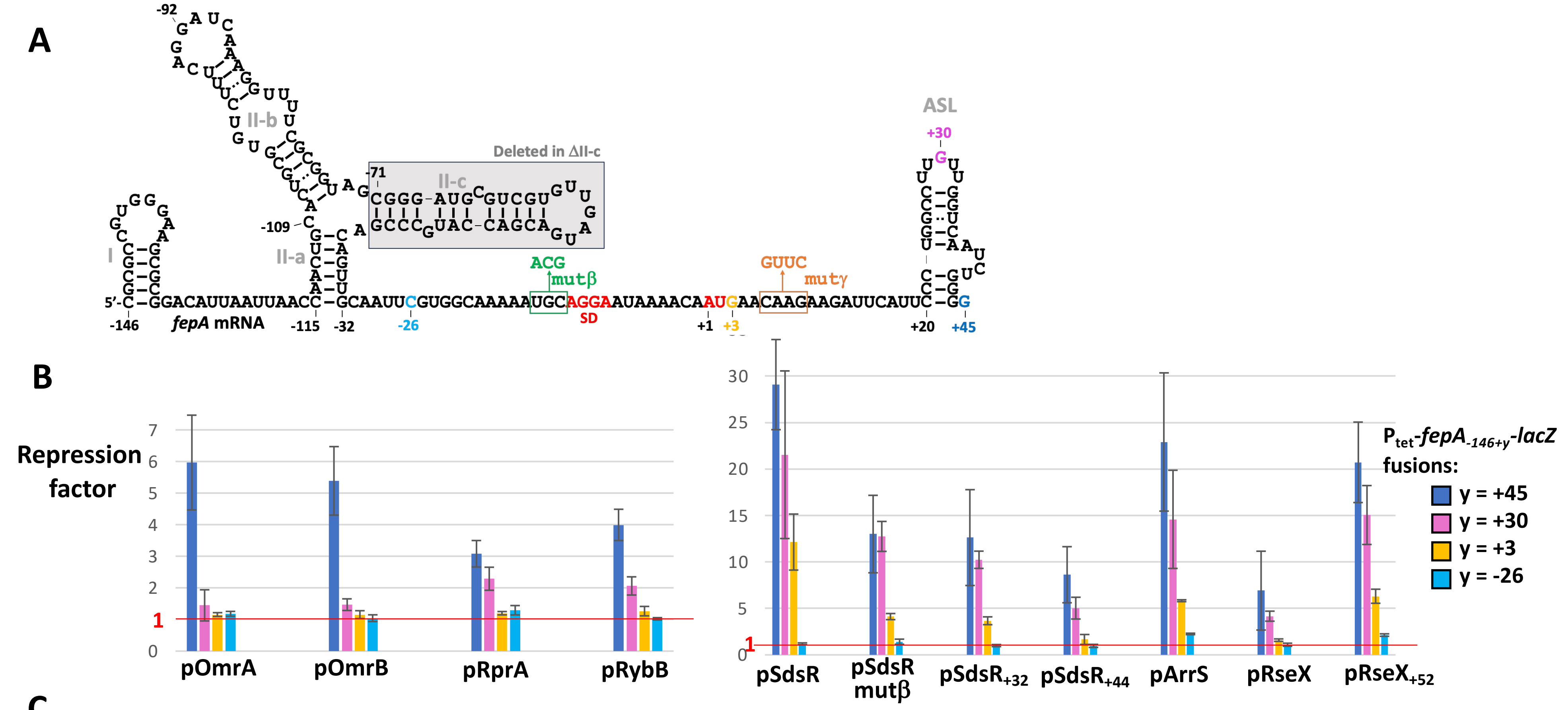

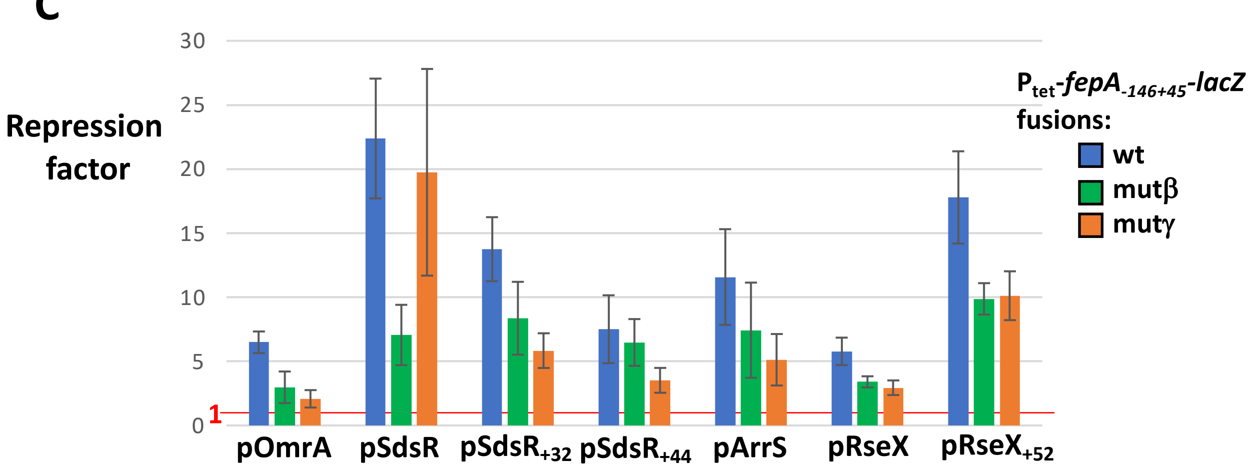

**Fig. S4. *fepA* translation initiation region is required for control by the far5'UTR-dependent sRNAs.** The repression of the $P_{tet}$-*fepA*$_{-146+45}$-*lacZ fusion*, wt or mutated in the TIR was measured for the different sRNAs repressing *fepA*. Mutations used are depicted in (A). The results with fusions truncated on the 3' end of *fepA* are shown in (B), and with nt changes in the TIR in (C). Strains used in panel (B) are JJ389, ESF87, OK818 and ESF17, and in panel (C) JJ389, JJ330 and ES148. For the $P_{tet}$-*fepA*$_{-146-26}$-*lacZ* fusion, the *lacZ* TIR is present.

Fig. S4

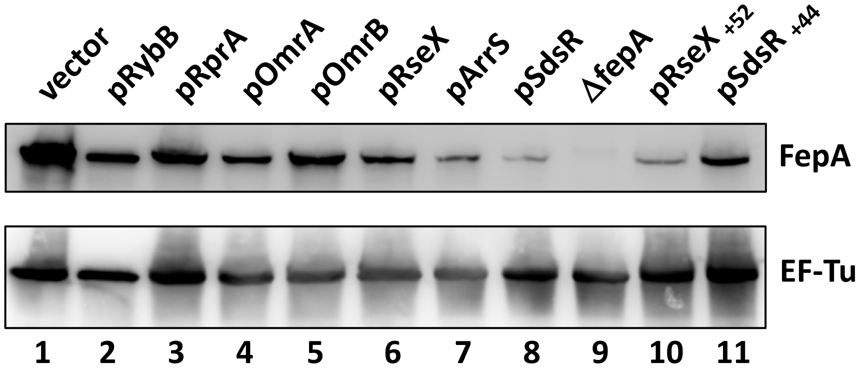

**Fig. S5. Western blot analysis of the FepA levels in the presence of plasmids overexpressing the indicated sRNAs.** The EF-Tu protein was detected from the same membrane and used as a loading control. Lanes 1 to 9 are presented in the Fig. 1C, and lanes 1 and 9-11 in Fig. 6E.

Fig. S5

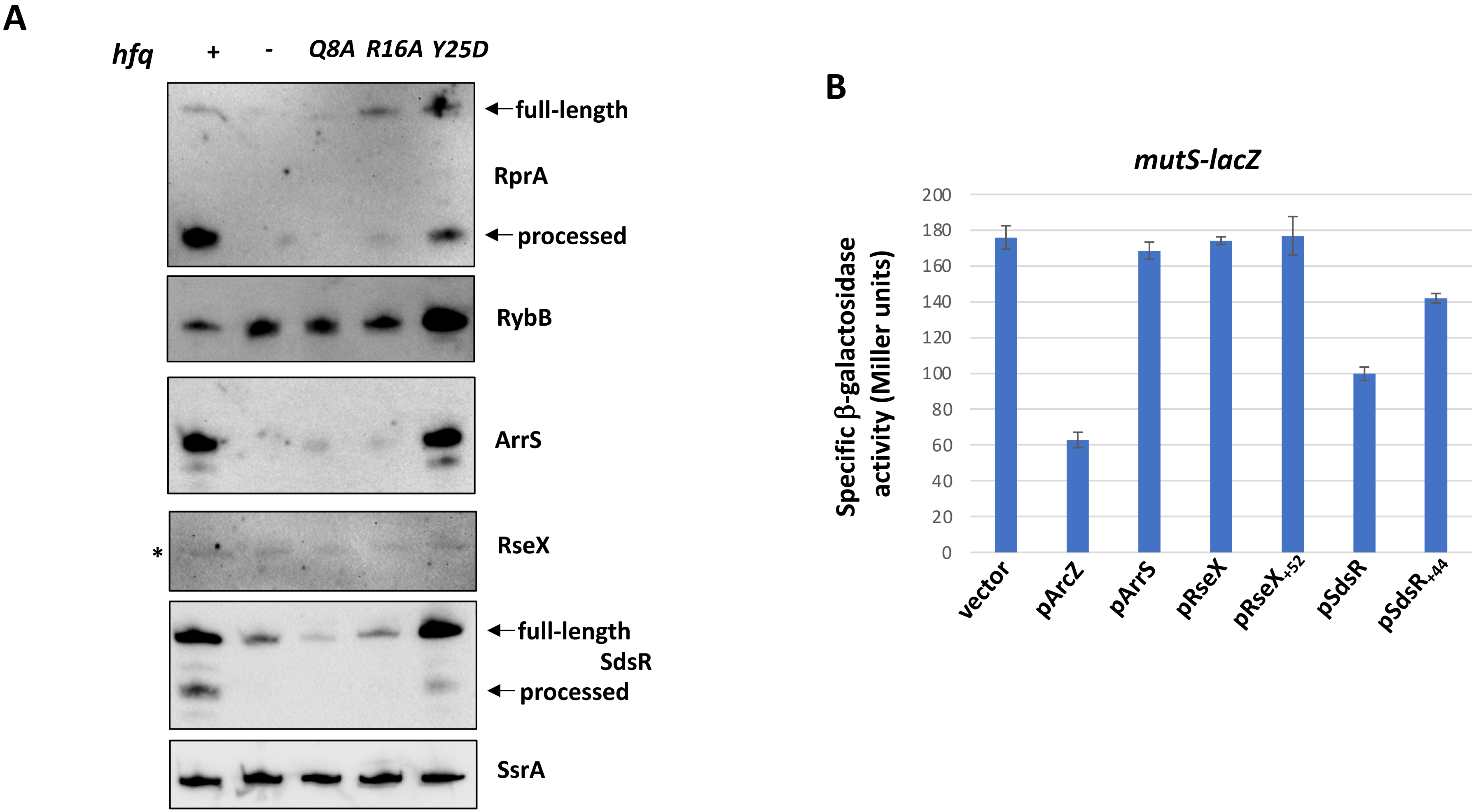

**Fig. S6. Hfq and the sRNAs repressing *fepA*.** (A) The levels of endogenous RprA, RybB, ArrS, RseX and SdsR sRNAs were followed by Northern blot in *hfq*$^{+}$, *Δhfq* strains and in *hfq* mutants of the proximal face (Q8A), rim (R16A) or distal face (Y25D). Cells were diluted 500-fold from overnight cultures, and growth was continued for 90 minutes after the cells reached an $OD_{600}$ of 2.4, before the RNA was extracted. The asterisk indicates an aspecific band detected with the RseX-specific probe (see also Fig. 2C). Strains used in this experiment are MG2412, MG2413, MG2414, MG2415 and MG2416. (B) The far5'UTR-dependent sRNAs repressing *fepA* do not affect other known Hfq-targets such as *mutS*. The β-galactosidase activity of the *mutS-lacZ* fusion in strain JC1060 (1) was measured upon overproduction of the indicated sRNAs. ArcZ was included as a positive control as it is a known repressor of *mutS* (1); SdsR also was previously shown to repress this fusion even if it does not carry a previously demonstrated pairing site (1, 7).

**Table S1. Strains and plasmids used in this study.**

| Strain name | Relevant characteristics | Source |
|---|---|---|
| MG1655 | E. coli wt reference strain for this study | From F. Blattner's lab |
| DJ480 | MG1655 *ΔlacX74* | D. Jin, NIH |
| DJ624 | MG1655 *ΔlacX74 mal::*$lacI^{q}$ | D. Jin, NIH |
| NM300 | DJ480 *mini-λ-tet* | N. Majdalani, NIH |
| NM1200 | MG1655 *mini-λ-cat* | N. Majdalani, NIH |
| ES017 | MG1508 $P_{tet}$-*fepA*$_{-146-26}$-*lacZ*$_{-17}$ | This study, Recombineering in MG1508 |
| ES021 | JJ389 Δ*fur::cat* | This study, JJ389 + P1 (MG1056) |
| ES023 | JJ389 Δ*fur::cat* Δ*fepA::gnt* | This study, ES21 + P1 (JJ0057) |
| ES024 | MG1325 Δ*fepA::gnt* | This study, MG1325 + P1 (JJ0057) |
| ES087 | MG1508 $P_{tet}$-*fepA*$_{-146+30}$-*lacZ*$_{+28}$ | This study, Recombineering in MG1508 |
| ES119 | OK410 Δ*sdsR::kan* | This study, OK410 + P1 (JJ0139) |
| ES120 | OK410 Δ*sdsR::kan* Δ*omrAB::tet* | This study, ES119 + P1 (MG1462) |
| ES121 | MG1432 Δ*arrS::cat* | This study, Recombineering in MG1432 |
| ES123 | OK410 Δ*sdsR::kan* Δ*omrAB::tet* Δ*arrS::cat* | This study, ES120 + P1 (ES121) |
| ES129 | OK410 Δ*sdsR::kan* Δ*omrAB::tet* Δ*arrS::cat* Δ*rseX::spc* | This study, ES123 + P1 (JJ156) |
| ES148 | MG1508 $P_{tet}$-*fepA*$_{-146+45}$mutγ-*lacZ*$_{+28}$ | This study, Recombineering in MG1508 |
| ES227 | ES129 D*arrS::FRT* (chloramphenicol-sensitive) | This study; cat cassette eliminated in ES129 |
| ES229 | MG1432 Δ*rprA::cat* | This study, Recombineering in MG1432 |
| ES230 | OK410 Δ*sdsR::kan* Δ*omrAB::tet* Δ*arrS::FRT* Δ*rseX::spc* Δ*rprA*::cat | This study, ES227 + P1 (ES229) |
| ES232 | ES230 D*rprA::FRT* (chloramphenicol-sensitive) | This study; cat cassette eliminated in ES230 |
| ES234 | OK410 Δ*hfq*::cat | This study, OK410 + P1 (MG1110) |
| ES250 | OK410 Δ*sdsR::kan* Δ*omrAB::tet* Δ*arrS::FRT* Δ*rseX::spc* Δ*rprA::FRT* Δ*rybB::cat-sacB* | This study, ES232 + P1 (KMT198) |
| KMT198 | Δ*rybB::cat-sacB* | K. Thompson/ S. Gottesman's lab, NIH |
| FQ99 | MG1508 $P_{tet}$-fepA$_{-146+45}$muta-lacZ$_{+28}$ | This study, Recombineering in MG1508 |
| FQ101 | MG1508 $P_{tet}$-fepA$_{-39+45}$-lacZ$_{+28}$ | This study, Recombineering in MG1508 |
| FQ125 | DJ624 $P_{tet}$-*fepA*$_{-146+45}$-*mScarlet (KanS)* | This study; *nptII* (KanR) eliminated in OK493 |
| FQ128 | DJ624 $P_{tet}$-*fepA*$_{-146+45}$-*mScarlet* | This study; OK523 + P1 (FQ125) |
| JC1060 | MG1655 *mal::*$lacI^{Q}$ *ΔaraBAD araC+ lacI'::kan-CP12b-mutS-lacZ* | (1) |

| | | |
|---|---|---|
| JJ0057 | MG1432 Δ*fepA::gnt* | This study, Recombineering in MG1432 |
| JJ0065 | DJ624 Δ*fepA::gnt* Δ*cirA::spc* Δ*fecA::catsacB* | Lab stock |
| JJ0066 | DJ624 Δ*fepA::gnt* Δ*cirA::spc* Δ*fecA::catsacB* Δ*omrAB::tet* | This study, JJ0065 + P1 (MG1462) |
| JJ0135 | MG1508 $P_{tet}$-*fepA*$_{-26+45}$-*lacZ*$_{+28}$ | (2) |
| JJ0136 | MG1508 $P_{tet}$-*fepA*$_{-92+45}$-*lacZ*$_{+28}$ | This study, Recombineering in MG1508 |
| JJ0139 | NM300 Δ*sdsR::kan* | This study, Recombineering in NM300 |
| JJ0146 | MG1508 $P_{tet}$-*fepA*$_{-26+45}$mutβ-*lacZ*$_{+28}$ | This study, Recombineering in MG1508 |
| JJ0156 | NM300 Δ*rseX::spc* | This study, Recombineering in NM300 |
| JJ0171 | DJ624 Δ*fur::cat* Δ*fepA::gnt* | This study, MG1169 + P1 (JJ0057) |
| JJ0184 | DJ624 Δ*fur::cat* Δ*omrAB::tet* | This study, MG1169 + P1 (JJ0066) |
| JJ0187 | DJ624 Δ*fur::cat* Δ*omrAB::tet* Δ*sdsR::kan* | This study, JJ0184 + P1 (JJ0139) |
| JJ0188 | DJ624 Δ*fur::cat* Δ*omrAB::tet* Δ*sdsR::kan* Δ*rseX::spc* | This study, JJ0187 + P1 (JJ0156) |
| JJ0199 | DJ624 Δ*fur::cat* Δ*omrAB::tet* Δ*sdsR::kan* Δ*rseX::spc* Δ*fepA::gnt* | This study, JJ0188 + P1 (JJ0057) |
| JJ0330 | MG1508 $P_{tet}$-*fepA*$_{-146+45}$mutβ-*lacZ*$_{+28}$ | Recombineering in MG1508 |
| JJ0389 | MG1508 $P_{tet}$-*fepA*$_{-146+45}$-*lacZ*$_{+28}$ | Recombineering in MG1508 |
| JJ0406 | MG1508 $P_{tet}$-*fepA*$_{-120+45}$-*lacZ*$_{+28}$ | Recombineering in MG1508 |
| JJ0407 | MG1508 $P_{tet}$-*fepA*$_{-109+45}$-*lacZ*$_{+28}$ | Recombineering in MG1508 |
| JJ0424 | MG1508 $P_{tet}$-*fepA*$_{-146+45}$loop1-*lacZ*$_{+28}$ | Recombineering in MG1508 |
| MG1056 | DJ480 Δ*fur::cat* | Lab stock, Δ*fur::cat* allele from (3) |
| MG1110 | DJ624 *hfq::cat* | (4) |
| MG1169 | DJ624 Δ*fur::cat* | Lab stock, Δ*fur::cat* allele from (3) |
| MG1325 | MG1099 *zce-726::Tn10* | (2) |
| MG1326 | MG1099 *rne-3071 zce-726::Tn10* | (2) |
| MG1432 | DJ624 *mini-λ-Tet* | Lab stock |
| MG1462 | NM1200 Δ*omrAB::tet* | This study, Recombineering in NM1200 |
| MG1508 | MG1655 *mal::*$lacI^q$*, mini-λ-Tet, mhpR*-$P_{tet}$-*cat-sacB-lacZ* | (5) |
| MG2379 | MG1508 $P_{tet}$-*fepA*$_{-146+45}$Stem5'-*lacZ*$_{+28}$ | This study, Recombineering in MG1508 |
| MG2381 | MG1508 $P_{tet}$-*fepA*$_{-146+45}$Stem3'-*lacZ*$_{+28}$ | This study, Recombineering in MG1508 |
| MG2382 | MG1508 $P_{tet}$-*fepA*$_{-146+45}$StemComp-*lacZ*$_{+28}$ | This study, Recombineering in MG1508 |

| | | |
|---|---|---|
| MG2384 | MG1508 $P_{tet}$-*fepA*$_{-146+45}$DII-c-*lacZ*$_{+28}$ | This study, Recombineering in MG1508 |
| MG2385 | MG1508 $P_{tet}$-*fepA*$_{-146+45}$loop2-*lacZ*$_{+28}$ | This study, Recombineering in MG1508 |
| MG2412 | DJ624 *hfq* wt $P_{tet}$-*fepA-mSc* (KanS) | This study, OK581+ P1 (FQ125) |
| MG2413 | DJ624 *Δhfq* $P_{tet}$-*fepA-mSc* (KanS) | This study, OK582+ P1 (FQ125) |
| MG2414 | DJ624 *hfqQ8A* $P_{tet}$-*fepA-mSc* (KanS) | This study, OK583+ P1 (FQ125) |
| MG2415 | DJ624 *hfqR16A* $P_{tet}$-*fepA-mSc* (KanS) | This study, OK584+ P1 (FQ125) |
| MG2416 | DJ624 *hfqY25D* $P_{tet}$-*fepA-mSc* (KanS) | This study, OK585+ P1 (FQ125) |
| OK410 | MG1508 low$P_{tet}$-*fepA*$_{-146+45}$-*lacZ* | This study, Recombineering in MG1508 |
| 0K493 | DJ624 $P_{tet}$-*fepA*$_{-146+45}$-*mScarlet* - FRT-*nptII*-FRT | This study, Recombineering in MG2352 |
| OK523 | DJ624 *ΔargG*::FRT-*nptII*-FRT | (6) |
| OK581 | DJ624 *ΔargG::FRT hfq*$^{+}$ | (6) |
| OK582 | DJ624 *ΔargG::FRT Δhfq* | (6) |
| OK583 | DJ624 *ΔargG::FRT hfqQ8A* | (6) |
| OK584 | DJ624 *ΔargG::FRT hfqR16A* | (6) |
| OK585 | DJ624 *ΔargG::FRT hfqY25D* | (6) |
| OK818 | MG1508 $P_{tet}$-*fepA*$_{-146+3}$-*lacZ*$_{+4}$ | This study, Recombineering in MG1508 |
| **Plasmid name** | **Characteristics** | **Reference** |
| pBRplac | $P_{LlacO-1}$ in pBR322, AmpR, TetR | (3) |
| pRprAmutα | rprAmutα under $P_{LlacO-1}$ in pBRplac, AmpR, TetR | This study, Mutagenic PCR |
| pMcaS | *mcaS* under $P_{LlacO-1}$ in pBRplac, AmpR, TetR | This study, NEB HiFi assembly |
| pRaiZ | *raiZ* under $P_{LlacO-1}$ in pBRplac, AmpR, TetR | This study, NEB HiFi assembly |
| pArrS | *arrS* under $P_{LlacO-1}$ in pBRplac, AmpR, TetR | This study, NEB HiFi assembly |
| pNarS (lab name : pNarS$_{+7}$) | *narS* under $P_{LlacO-1}$ in pBRplac, AmpR, TetR | This study, NEB HiFi assembly, this plasmid differs from pNarS in (4) because it contains only 7 nts after the Us stretch of the terminator |
| pCsrC | *csrC* under $P_{LlacO-1}$ in pBRplac, AmpR, TetR | This study, NEB HiFi assembly |
| pSgrS | *sgrS* under $P_{LlacO-1}$ in pBRplac, AmpR, TetR | This study, NEB HiFi assembly |
| pSdsRmutβ | sdsRmutβ under $P_{LlacO-1}$ in pBRplac, AmpR, TetR | This study, Mutagenic PCR |
| pSdsR$_{+32}$ | *sdsR*$_{+32}$ under $P_{LlacO-1}$ in pBRplac, AmpR, TetR | This study, Mutagenic PCR |
| pSdsR$_{+44}$ | *sdsR*$_{+44}$ under $P_{LlacO-1}$ in pBRplac, AmpR, TetR | This study, Mutagenic PCR |
| pSdsR$_{+44}$mut | *sdsR*$_{+44}$*mut* under $P_{LlacO-1}$ in pBRplac, AmpR, TetR | This study, Mutagenic PCR |
| pRseX$_{+52}$ | *rseX*$_{+52}$ under $P_{LlacO-1}$ in pBRplac, AmpR, TetR | This study, Mutagenic PCR |

| | | |
|---|---|---|
| pRseX+52mut | *rseX+52mut* under $P_{LlacO-1}$ in pBRplac, AmpR, TetR | This study, Mutagenic PCR |

**Table S2. Oligonucleotides used in this study**

| Name | Sequence | Use |
|---|---|---|
| ***Strains construction*** | | |
| 5'DomrAB ::tet | GCGAAACGCTGTTGCGATTGACCGCTGGTGGCGTTTGGCTT CAGGTTGC**TCCTAATTTTTGTTGACACTCTA** | Construction *ΔomrAB::tet* allele (strain MG1462) |
| 3'DomrAB ::tet | CGCGAGCGACAGTAAATTAGGTGCGAAAAAAAACCTGCGC ATCCGCGCAGGTT**CTCTTGGGTTATCAAGAGGG** | |
| 5'fepA::gnt | GCAATTCGTGGCAAAAATGCAGGGACGCACACCGTGGAAA CGG | Construction *ΔfepA::gnt* allele (strain JJ0057) |
| 3'fepA::gnt | GGTGTTTACGCTCATATACCACGGCGGCGTTGTGACAATTT ACC | |
| 5'RyeB::kanRfor | CGCAAACTGGAAAACCTGGCGTCGTCATCTATTCTTAAAGA AAGCCACGTTGTGTCTCAA | Construction *ΔsdsR::kan* allele (strain JJ0139) |
| 3'RyeB::kanRrev | GCGAAGGAAATGCTTCTGGCTTTTAACAGATAAAAAGAGA CCGCTGGAATTGGGAATTGATCC | |
| ESF134 | AATAAGTAATCCGGGTTCATTTTTTTGCAACTGGCGTTGAgt gtaggctggagctgcttcgaagt | Construction *ΔarrS::cat* allele (strain ES121) |
| ESF135 | CCAGTTTGTGATCTCTGAAGAATATTACTAAAGTTAAAATtc ctccttagttcctattccgaag | |
| ESF173 | GGTAGTACCTGTCGCAAATTCTTTACAGTTTTTAAACTAAgt gtaggctggagctgcttcgaagt | Construction *ΔrprA::cat* allele (strain ES229) |
| ESF174 | CGGCTTGAAGAGAGTCACAGTATCTTGTGCAACATTATTGtc ctccttagttcctattccgaag | |
| 5'forRseX::spc | GCTTTATTAATTCATTTAATCAATATATTAGCACTGATTAC AATTATACCAAACGGATGAAGGCACGAA | Construction *ΔrseX::spc* allele (strain JJ0156) |
| 3'revRseX::spc | CCAAACGGCTGGTGTGATCAGGCGCACATTAATGAAGGCA TTTATTTGCCGACTACCTTGG | |
| ***Strains construction : lacZ and mScarlet fusions*** | | |
| Ptet-55-12for | ctccctatcagtgatagagattgacatccctatcagtgatagag | Forward primer to increase homology region if necessary (in the Ptet region) |
| lacZ4-67rev | taacgccagggttttcccagtcacgacgttgtaaaacgacggccagtgaatccgtaatcatggt | Reverse primers to increase homology region if necessary (in the lacZ coding sequence) |
| lacZ28-66rev | aacgccagggttttcccagtcacgacgttgtaaaacgac | |
| 5'Ptet fepA-146 | GATAGAGATTGACATCCCTATCAGTGATAGAGATACTGAG CACCGCGCCGTGGGAAGCGCGG | Forward primer for construction of $P_{tet}$-fepA$_{-146}$ … fusions |
| 3'fepA-l-lac | TAACGCCAGGGTTTTCCCAGTCACGACGTTGTAAAACGACC CCCAGATTGACCAACAAGG | Reverse primer for construction of *fepA*$_{+45}$-*lacZ* fusions |
| AK08b | CATTCGCCATTCAGGCTGCGCAAC | Primers for low$P_{tet}$-*fepA*$_{-146+45}$-*lacZ* fusion (strain OK410) |
| AK283 | TGACACCATCGAATGGCGCTCCCTATCAGTGATAGAGATG GACATCCCTATC | |
| AK386 | AACGCATAAATTCTTTTATTACTGCTTCGCCTTTGCTCACCC CCAGATTGACCAACAAGG | Reverse primer for $P_{tet}$-*fepA*$_{-146+45}$-*mScarlet* fusion (strain OK493), used with 5'Ptet fepA-146 |
| ESF20 | AGTGAATCCGTAATCATGGTCATAGCTGTTTCCTGTGTGAA ATTGCAACTGTCGGGCATG | Reverse primer for $P_{tet}$-*fepA*$_{-146-26}$-*lacZ*$_{-17}$ fusion (strain ES17), used with 5'Ptet fepA-146 and lacZ4-67rev |
| ESF98 | TCACGACGTTGTAAAACGACCAAGGCCAGGGAATGAATCT TCTTGTTCATTGTTTTATTCCTGCATTTTT | Reverse primer for $P_{tet}$-*fepA*$_{-146+30}$-*lacZ*$_{+28}$ fusion (strain ES87), |

| | | |
|---|---|---|
| | | used with 5'Ptet fepA-146 |
| FQP70 | CCATGCCCGACAGTTGCAAAAGCTGGCAAAAATGCAGGAATAAAAC | Primers for $P_{tet}$-*fepA*$_{-146+45}$mutα-*lacZ*$_{+28}$ fusion (strain FQ99), used with 5'Ptet fepA-146 and 3'fepA-l-lac |
| FQP71 | GTTTTATTCCTGCATTTTTGCCAGCTTTTGCAACTGTCGGGCATGG | |
| FQP74 | CATCCCTATCAGTGATAGAGATACTGAGCACGACAGTTGCAATTCGTGGCAAAAATG | Forward primer for $P_{tet}$-*fepA*$_{-39+45}$-*lacZ*$_{+28}$ fusion (strain FQ101), used with 3'fepA-l-lac |
| 5-PtetFepA-92 | GATAGAGATTGACATCCCTATCAGTGATAGAGATACTGAGCACGATCAAAGGTTTTCGCGGTAGC | Forward primer for $P_{tet}$-*fepA*$_{-92+45}$-*lacZ*$_{+28}$ fusion (strain JJ0136), used with 3'fepA-l-lac |
| 5PtetFepA-26mutRB | GATAGAGATTGACATCCCTATCAGTGATAGAGATACTGAGCACCGTGGCAAAAAACGAGGAATAAAAC | Forward primer for $P_{tet}$-*fepA*$_{-26+45}$mutβ-*lacZ*$_{+28}$ fusion (strain JJ0146), used with 3'fepA-l-lac |
| 5'fepAmutRBfor | CGTGGCAAAAAACGAGGAATAAAACAatgAACAAGAAG | Primers for $P_{tet}$-*fepA*$_{-146+45}$mutβ-*lacZ*$_{+28}$ fusion (strain JJ0330), used with 5'Ptet fepA-146 and 3'fepA-l-lac |
| 3'fepAmutRBrev | GTTTTATTCCTCGTTTTTTGCCACGAATTGCAACTGTCG | |
| 5'PtetfepA-120 | CCCTATCAGTGATAGAGATACTGAGCACTTAACCAACTGCACTGCGTGTCTTTCAGG | Forward primer for $P_{tet}$-*fepA*$_{-120+45}$-*lacZ*$_{+28}$ fusion (strain JJ0406), used with 3'fepA-l-lac |
| 5'PtetfepA-109 | CCCTATCAGTGATAGAGATACTGAGCACCACTGCGTGTCTTTCAGGATCAAAGG | Forward primer for $P_{tet}$-*fepA*$_{-109+45}$-*lacZ*$_{+28}$ fusion (strain JJ0407), used with 3'fepA-l-lac |
| 5'fepAmutRX3 | GCGTGTCTTTGTCGATCAAAGGTTTTCGCGGTAGC | Primers for $P_{tet}$-*fepA*$_{-146+45}$loop1-*lacZ*$_{+28}$ fusion (strain JJ0424), used with 5'Ptet fepA-146 and 3'fepA-l-lac |
| 3'fepAmutRX3 | CCTTTGATCGACAAAGACACGCAGTGCAGTTGG | |
| 5'fepAmutRX2for | CAGGATCATTCCTTTTCGCGGTAGCGGGATGC | Primers for $P_{tet}$-*fepA*$_{-146+45}$Stem3'-*lacZ*$_{+28}$ fusion (strain MG2381), used with 5'Ptet fepA-146 and 3'fepA-l-lac |
| 3'fepAmutRX2rev | CCGCGAAAAGGAATGATCCTGAAAGACACGCAG | |
| fepAmutRX2Lfor | cactgcgtgtGGAAcaggatcaaaggttttcgcgg | Primers for $P_{tet}$-*fepA*$_{-146+45}$Stem5'-*lacZ*$_{+28}$ fusion (strain MG2379), used with 5'Ptet fepA-146 and 3'fepA-l-lac |
| fepAmutRX2Lrev | ctttgatcctgTTCCacacgcagtgcagttggttaattaatg | |
| fepARX2L+Rfor | cactgcgtgtGGAAcaggatcaTTCCttttcgcgg | Primers for $P_{tet}$-*fepA*$_{-146+45}$StemComp-*lacZ*$_{+28}$ fusion (strain MG2382), used with 5'Ptet fepA-146 and 3'fepA-l-lac. Note: PCR amplification was done on a template carrying the Stem3' mutation |
| fepARX2L+Rrev | GAAtgatcctgTTCCacacgcagtgcagttggttaattaatg | |
| fepAmutRX62for | GTGTCTTTCAGGAagtAAGGTTTTCGCGGTAGCGGG | Primers for $P_{tet}$-*fepA*$_{-146+45}$loop2-*lacZ*$_{+28}$ fusion (strain MG2385), used with 5'Ptet fepA-146 and 3'fepA-l-lac |
| fepAmutRX62rev | GCGAAAACCTTactTCCTGAAAGACACGCAGTGCAG | |

| | | |
|---|---|---|
| fepAD-71-39for | GTTTTCGCGGTAGACAGTTGCAATTCGTGGCAAAAATG | Primers for $P_{tet}$-*fepA*$_{-146+45}$ΔII-c-*lacZ*$_{+28}$ fusion (strain MG2384), used with 5'Ptet fepA-146 and 3'fepA-l-lac (Position of the deleted region is between the two underlined nts) |
| fepAD-71-39rev | GAATTGCAACTGTCTACCGCGAAAACCTTTGATCCTG | |
| fepAmutArrS2lacZR | GTAAAACGACCCCCAGATTGACCAACAAGGCCAGGGAATGAATCTTGAACTTCATTGTTTATTCCTGCATTTTT | Reverse primer for $P_{tet}$-*fepA*$_{-146+45}$mutγ-*lacZ*$_{+28}$ fusion (strain ES148), used with 5'Ptet fepA-146 and 3'fepA-l-lac |
| AK671 | GGCAAAAATGCAGGAATAAAACAATGACCATGATTACGGATTCAC | Forward primer for $P_{tet}$-*fepA*$_{-146+3}$-*lacZ*$_{+4}$ fusion (strain OK818): used with primer lacZ28-66rev to amplify lacZ region and, in a 2nd PCR reaction, PCR#1 product was used with primer Ptet-55-12for to amplify the Ptet-fepA region (from JJ389 genomic DNA) |
| ***Plasmids construction*** | | |
| FQP72 | GAAATCCCCTGAGTGAAACAAGCTTTTGCTGTGTGTAGTCTTTG | Construction pRprAmutα |
| FQP73 | CAAAGACTACACACAGCAAAAGCTTGTTTCACTCAGGGGATTTC | |
| FQP122 | gagcggataacaagatactgacgtcATTGATCAACAAGCTGGAACGGCAG | Construction pRaiZ |
| FQP123 | taagctgtcaaacatgagaattcACTGTTTTCACGCTGTCAACAAAAAACGC | |
| placAatNarsF | gagcggataacaagatactgacgtcTCTATATCGCCTGCGTAGTGATTAC | Construction pNarS$_{+7}$ |
| FQP57 | gataagctgtcaaacatgagaattcGACGCATAAAAAAAGCGCGACATCATGC | |
| placAatrev (ESF1) | GACGTCAGTATCTTGTTATCCGCTC | Construction pBRplac derivatives (amplification of the vector backbone) |
| placEcofor (ESF2) | gaattctcatgtttgacagcttatc | |
| placAatSdsR+32 (ESF3) | GAGCGGATAACAAGATACTGACGTCCTTTTAGCGCACGGCTCTCTCC | Construction pSdsR$_{+32}$ |
| placEcoSdsRrev (ESF4) | gataagctgtcaaacatgagaattcGCGAAGGAAATGCTTCTGGCTT | |
| 5'RyeBmutfor | GGCAAGGCAACTAAGCCTCGTTTAATGCCAACTTTTAGCGCACG | Construction pSdsRmutβ |
| 3'RyeBmutrev | CGTGCGCTAAAAGTTGGCATTAAACGAGGCTTAGTTGCCTTGCC | |
| SdsR+44F (ESF41) | GATAACAAGATACTGACGTCGGCTCTCTCCCAAGAGCCATTTCCC | Construction pSdsR$_{+44}$ |
| SdsR+44R (ESF42) | GGGAAATGGCTCTTGGGAGAGAGCCGACGTCAGTATCTTGTTATC | |

| | | |
|---|---|---|
| SdsR+44mut1for (ESF187) | GCCATTTCCCTGGACCGAATACAccttTCGTGTTCGGTCTCTTTTTATCTG | Construction pSdsR$_{+44}$mut |
| SdsR+44mut1rev (ESF188) | CAGATAAAAAGAGACCGAACACGAaaggTGTATTCGGTCCAGGGAAATGGC | |
| RseX+52F (ESF45) | AGCGGATAACAAGATACTGACGTCGTTTATATTTTTGGGCCGGC | Construction pRseX$_{+52}$ |
| RseX+52R (ESF46) | GCCGGCCCAAAAAATATAAACGACGTCAGTATCTTGTTATCCGCT | |
| RseX+52mut6for (ESF185) | gtcGTTTATATTTTTGGGCCGGCAactTGCCGGCTTTTTtttATGCCTTC | Construction pRseX$_{+52}$mut |
| RseX+52mut6rev (ESF186) | GAAGGCATaaaAAAAAGCCGGCAagtTGCCGGCCCAAAAATATAAACgac | |
| PlacAatSgrSF (ESF72) | GAGCGGATAACAAGATACTGACGTCGATGAAGCAAGGGGGTGCCCCATGC | Construction pSgrS |
| PlacEcoSgrSR (ESF73) | GATAAGCTGTCAAACATGAGAATTCCATACGTTCCCTTTTTAGCGCGGCG | |
| PlacAatArrSF (ESF82) | GAGCGGATAACAAGATACTGACGTCGTAATCCGATTTAAATATCGAGTCTC | Construction pArrS |
| PlacEcoArrSR (ESF83) | GATAAGCTGTCAAACATGAGAATTCCAGGTAATTTCAATGCGTTAAAAG | |
| PlacAatCsrCF (ESF61) | GAGCGGATAACAAGATACTGACGTCATAGAGCGAGGACGCTAACAGGAAC | Construction pCsrC |
| PlacEcoCsrCR (ESF62) | GATAAGCTGTCAAACATGAGAATTCTTTGCGGCGGAATCTAACAGAAAGC | |
| PlacAatMcaSF (ESF74) | GAGCGGATAACAAGATACTGACGTCACCGGCGCAGAGGAGACAATGC | Construction pMcaS |
| PlacEcoMcaSR (ESF75) | GATAAGCTGTCAAACATGAGAATTCGCGGCTATCTGCAAAGTTAAAACTGC | |
| ***Templates for T7 in vitro transcription*** | | |
| pT7fepA | TAATACGACTCACTATAGGCTATTTGCATTTGCAATAGCG | Forward primer for template for *in vitro* transcription of *fepA* (from-173) |
| 5'-pT7-fepA-146+3G | TAATACGACTCACTATAGGGCGCGCCGTGGGAAGCGCGGAC | Forward primer for template for *in vitro* transcription of *fepA* (from-146) |
| seqfepA99rev | CTGCTCGGCGGCGGTAACGACAATAGTATCG | Reverse primer for template for *in vitro* transcription of *fepA* (up to +129) |

| | | |
|---|---|---|
| 3’fepA26-45rev | CCCCAGATTGACCAACAAGG | Reverse primer for template for *in vitro* transcription of *fepA* (up to +45) |
| 5-pT7RyeB | TAATACGACTCACTATAggcAAGGCAACTAAGCCTGC | Forward and reverse primers for template for *in vitro* transcription of SdsR wt and SdsRmutβ |
| 3-RyeB+81rev | AAAAAGAGACCGAACACGATTCC | |
| 5-pT7-SdsR44 | TAATACGACTCACTATAGGCTCTCTCCCAAGAGCCATTTCCC | Forward primer for template for *in vitro* transcription of SdsR$_{+44}$ (with primer 3-RyeB+81rev) and SdsR$_{+44}$mut (with primer 3-SdsRmutTER+81rev) |
| 3-SdsRmutTER+81rev | AAAAAGAGACCGAACACGAAAGG | Reverse primer for template for *in vitro* transcription of SdsR$_{+44}$mut (with primer 5-pT7-SdsR44) |
| pT7G-RseX+1+52long | TAATACGACTCACTATAGTTTTATTATTCTGTGTCATGATGCTTCCGTTATTAGCCTTTTATCGTCTTG | Forward and reverse primers for template for *in vitro* transcription of RseX wt (primer annealing and PCR) |
| RseXrev+91+24long | AAAAAAAAGCCGGCATCATGCCGGCCCAAAAAATATAAACAAGACGATAAAAGGCTAATAACGGAAGC | |
| ***Cy5-oligonucleotides for reverse-transcription*** | | |
| FepAToeCy5 | Cy5-CGGTAACGACAATAGTATCG | RT primer used for DMS, CMCT and RNases probing |
| FepAToeCy5+26+45 | Cy5-CCCCAGATTGACCAACAAGG | RT primer used for Lead-acetate probing |
| FepAToeCy5 2L | Cy5-CGTCATGTGAAACAGGAGTATCGGTCGGCTCT | Used for reverse-transcription from total RNA |
| ***Biotinylated probes for Northern-blots (5’-BioTEG)*** | | |
| NarS-probe | GCAGCACATGTAACCCGAAGTATGACGAGTAACGGTTTATTTTTTAG | |
| RaiZ-probe | CGATACACTCAATATAAAGGActaCTCTTCTTCAACTTCTTCG | |
| CsrC-probe | GGAACAATCCGTGTTGATTCCATTTCCGTTTAATTACGTC | |
| SdsRterm-Probe (ESF24) | AAAAAGAGACCGAACACGATTCCTGTATTCGGTCCA | |
| RseXterm-Probe (ESF25) | AAAAAAAAGCCGGCATCATGCCGGCCCAAAAAATATAAA | |
| RseX+52mut6-probe (ESF179) | AAAAAAAAGCCGGCAAGTTGCCGGCCCAAAAAATATAAAC | Complementary to the RseX$_{+52}$mut sequence |
| ArrS-probe (ESF139) | CCAGCTTAAGTCGAAACAAGGAGACTCGATATTTAAATCG | |
| SgrS-probe (ESF142) | AACGCAACCAGCACAACTTCGCTGTCGCGGTAAAATAGTG | |
| RprA-probe | CAGGGGATTTCCATGCTTATAAATCAATATGTTGATTTATAACc | |

| | |
|---|---|
| RybB-probe | GCTCCACAAAATGGGGACATCAAAGAAAAGCAGTGGC |
| OmrA-probe | caggttggtgcaagagacagggtacgaagagcgtaccg |
| OmrB-probe | cgcaggctggtgtaattcatgtgctcaacccgaagttga |
| SsrA-probe | CGCCACTAACAAACTAGCCTGATTAAGTTTTAACGCTTCA |
| 5S-probe | CTACCATCGGCGCTACGGCGTTTCACTTCTGAGTTCG |
| ompA-probe | ccattgttgttgatgaaaccagtgtcatggtactgggaccagc |
| fepA-probe | CGGCTTGCCGTCAATCAAAATCAGCGTGTTTTCCGGACC |
